# Gene-by-environment interactions and adaptive body size variation in mice from the Americas

**DOI:** 10.1101/2024.10.03.616282

**Authors:** Katya L. Mack, Nico P. Landino, Mariia Tertyshnaia, Tiffany C. Longo, Sebastian A. Vera, Lilia A. Crew, Kristi McDonald, Megan Phifer-Rixey

## Abstract

The relationship between genotype and phenotype is often mediated by the environment. Moreover, gene-by-environment (GxE) interactions can contribute to variation in phenotypes and, in turn, fitness. Nevertheless, understanding the impact of GxE interactions in wild systems remains challenging. In the last 500 years, house mice have invaded the Americas. Despite their short residence time, there is evidence of rapid climate adaptation, including shifts in body size and aspects of metabolism with latitude. Previous studies in this system have identified candidate genes for metabolic adaptation using selection scans, however, environmental variation in diet as well as GxE interactions affecting metabolism are likely important factors in shaping body mass variation in wild populations. Here, we investigate the role of the environment and GxE interactions in shaping adaptive phenotypic variation with an experimental manipulation of diet. Using new locally adapted inbred strains from North and South America, we evaluated response to a high-fat diet, finding that sex, strain, diet, and the interaction between strain and diet contribute significantly to variation in aspects of body size. We also found that transcriptional response to diet is largely strain-specific, indicating that GxE interactions affecting gene expression are pervasive. Next, we used crosses between strains from contrasting climates (New York x Brazil and New York x Florida) to characterize gene expression regulatory divergence on a standard breeder diet and on a high-fat diet. We found that gene regulatory divergence is often condition-specific, particularly for *trans*-acting changes. Finally, we find evidence for lineage-specific selection on *cis*-regulatory variation involved in diverse processes, including lipid metabolism. Overlap with scans for selection identified candidate genes for environmental adaptation with diet-specific effects. Together, our results underscore the importance of environmental variation and GxE interactions to adaptive variation in complex traits.

## Introduction

Understanding how populations vary in genotype, phenotype, and fitness in association with the local environment is critical to understanding the process of adaptation. One enduring challenge is integrating the simultaneous influence of the environment on the generation of and selection for phenotypic variation. While environmental factors can act as selective agents favoring particular phenotypes, they can also mediate the relationship between genotype and phenotype. A single genotype may produce different phenotypes across different environments, known as phenotypic plasticity [1–3]. These plastic responses can allow individuals to rapidly respond to environmental variability [4,5]. However, phenotypic plasticity itself can vary substantially among individuals or populations and be the target of selection [6,7]. When the environment influences the relationship between genotype and phenotype, this is known as a genotype-by-environment (GxE) interaction [1,8]. GxE interactions can result in phenotypic differences between individuals that are only observed under specific conditions, making them difficult to study in many natural populations [3]. However, the context-dependent nature of GxE interactions also means these interactions can play an important role in shaping the relationship between environment and organismal fitness during local adaptation, particularly when GxE interactions impact the expression of adaptive phenotypes [9–14].

House mice (*Mus musculus domesticus*) have proven to be a valuable system for studying local adaptation. Alongside humans, in the last ∼500 years, house mice have colonized a wide range of habitats and climates in the Americas [15,16]. Populations across latitudinal gradients show significant evidence of local adaptation at several complex traits [17–20]. One of the most striking examples is body mass. Past studies have identified variation in body size in house mice in association with latitude [17,18,21], consistent with the ecogeographic pattern known as Bergman’s rule [22]. In mammals, the covariance of body size and latitude is generally attributed to thermoregulatory adaptations to differences in temperature [23]. At higher latitudes, where individuals must contend with colder climates, larger body sizes can diminish heat loss through lower surface area to volume ratios. For example, mice from equatorial Manaus, Brazil, weigh nearly 37% less on average than mice from New York state [21]. These differences in body mass persist in the lab after multiple generations, indicating they have a genetic basis[18,24]. Genome-wide scans for selection and gene expression analyses have identified a number of candidate genes for climate adaptation, including genes with previously described roles in metabolism and body size variation [18–20,25–27]. Candidate variants for environmental adaptation identified so far are predominantly non-coding, suggesting that changes in gene expression regulation play an important role in driving adaptive body size evolution [18,26,28]. For example, *cis-*regulatory variation at genes *Adam17* and *Bcat2* was significantly associated with clinal body mass variation in wild mice in North America [26].

Less is known about the role of GxE interactions in metabolic and body size variation in this system. In humans and mice, genetics, diet, and gene-by-environment interactions have all been shown to contribute to variation in body mass and composition [29–31]. In humans, genotype-by-diet interactions play a major role in metabolic syndrome and obesity risk. A now well-described example comes from the fat mass and obesity-associated (FTO) locus [32].

Interactions between FTO risk alleles and diet (e.g., fatty foods) are associated with increased obesity risk [33]. The interaction between genetic and environmental factors related to diet can also play an important role in local adaptation. Variation in food abundance or nutritional value can exert strong selection pressure. For example, there is evidence that Greenlandic Inuits have adapted to lipid-rich diets [34]. Selection on genetic variants related to fat metabolism may have facilitated tolerance of diets high in omega-3 polyunsaturated fatty acids from seafood and variation in those loci is associated with differences in body weight and height. Similarly, differences in access to food resources may have exerted selection pressure as house mice colonized new climates in the Americas. Studies in wild mice indicate diet can vary drastically by location and season and that survival and breeding can be affected by food availability [35–39]. Consequently, climatic and geographic variation affecting diet may contribute to local adaptation in house mice. Previous studies have also suggested an important role for plastic responses in phenotypic and gene expression variation in this system [25,28,40]. For example, the temperature at which mice are reared has been found to affect aspects of morphology and gene expression variation[25,40]. When mice from Manaus, Brazil and Saratoga Springs, New York, were reared under cold temperatures, extremities (i.e., tail and ear length) were shorter than when mice were reared under warm conditions, consistent with Allen’s Rule [40]. Gene-by-temperature interactions affecting gene expression were also common (affecting 5-10% of genes surveyed) [25].

Here, we capitalize on new wild-derived inbred strains of house mice collected from North and South America to gain insight into the effects of genetic, environmental, and GxE interactions on body size and gene expression variation. First, we ask whether adaptive variation in body mass among populations is influenced by diet and genotype-by-diet interactions. Then, we use RNA-seq of liver tissue to investigate the influence of genetic and environmental variation on gene expression. Focusing on strains from divergent climates (New York and Florida, New York and Brazil), we identify gene regulatory differences between strains across diets as well as *cis*-by-diet and *trans*-by-diet interactions affecting expression divergence. Finally, we examine evidence for selection on gene regulatory changes to identify candidates for environmental adaptation. Overall, our results highlight the importance of the relationship between environmental and phenotypic variation during adaptive evolution.

## Results and Discussion

### Variation in phenotypic responses to a high-fat diet

Wild and lab-born mice from different populations in the Americas have been shown to differ in aspects of body size, largely consistent with Bergmann’s Rule [18,24]. Body size is a complex trait, dependent on many genes and the environment (e.g., [41,42]). Among environmental variables, diet is well-established as a factor that contributes significantly to variation in body weight (e.g., [42,43]). While house mice are human commensals, they are opportunistic omnivores and diet, estimated via carbon (δ^13^C) and nitrogen (δ^15^N) stable isotopes, has been shown to vary among wild populations in the Americas ([21], Kruskal–Wallis, δ^15^N *p*=7.96x10^-5^, δ^13^C *p*=6.60 x10^-4^; **Table S1, Fig. S1**). Moreover, populations at different latitudes are expected to differ in seasonal resource availability, making metabolic response to diet critical to fitness. We asked whether eight new wild derived strains from the Americas varied in aspects of body size and growth on a regularl breeder diet (hereafter, “regular” for simplicity) and a high-fat diet over twelve weeks post-weaning. Because house mice are sexually dimorphic (e.g., [44,45]), with male mice larger than female mice as seen in our data (**Figs. 1**, **S2**; **Tables S2-5**), we analyzed the data for each sex separately. As expected, we found the covariate in all analyses (either wean weight or week 12 body length), contributed significantly to variation in all measures (**Table 1**). We also found that strain contributed significantly to variation in all measures, with the exception of body length in females (**Table 1**). Tukey’s tests suggest that strains from higher latitude locations tend to be larger than those from lower latitude locations (**Tables S6, S7**). A notable exception is SARC, a strain from New York, USA, which is smaller than other high latitude strains. This result is not unexpected. Body size is not a fixed trait and varies within populations and each wild-derived strain represents a subset of variation sampled from a larger population. SARC showed similar results in a previous study, with body sizes closer to strains from lower latitudes and lower growth. Overall, differences among strains are largely consistent with results from wild populations and early generations of inbreeding [18,46] as well as previous phenotyping of rederived inbred strains [24].

**Figure 1.**
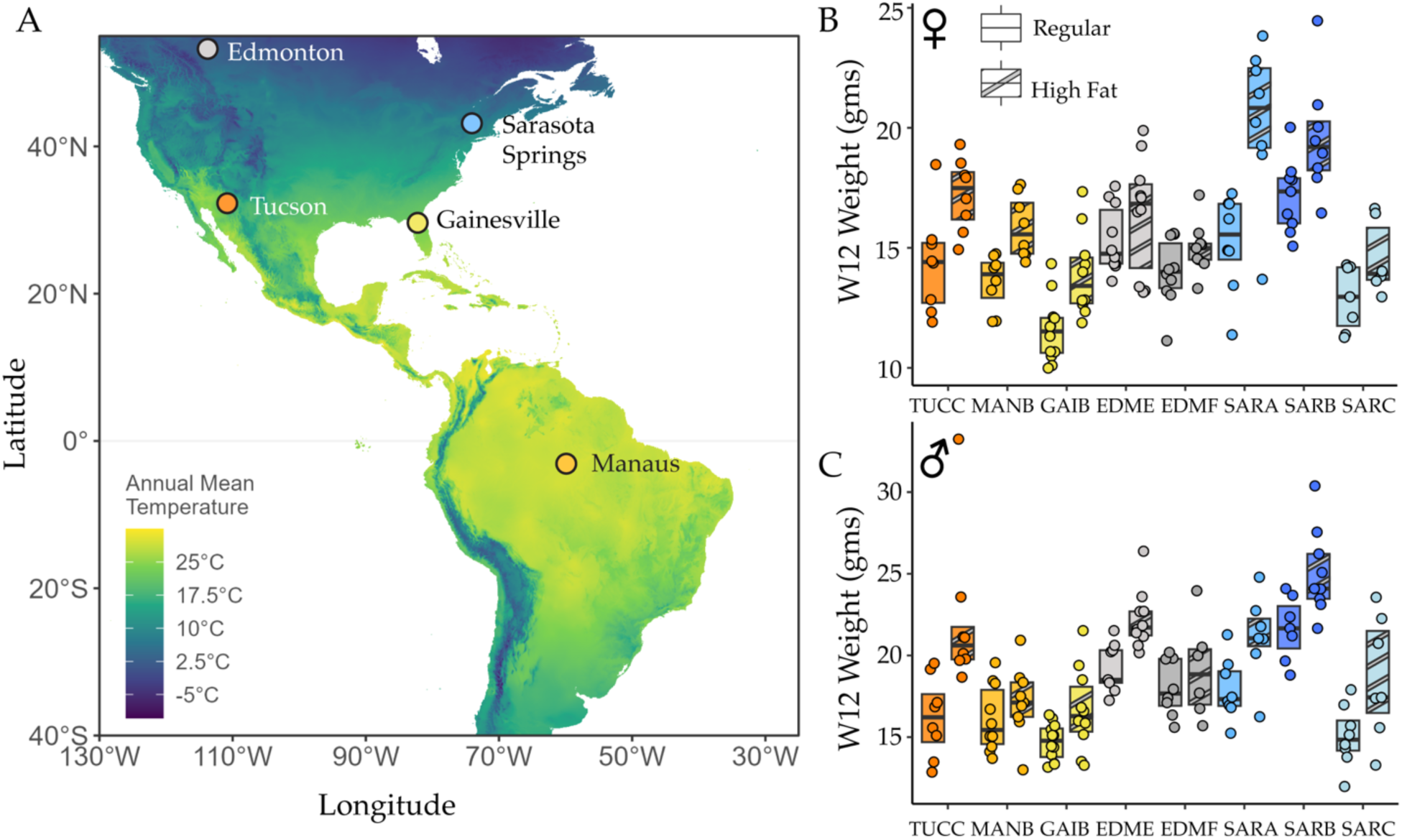
Variation in body size and response to diet among strains from the Americas. A) The new wild derived inbred strains originate from locations in the Americas. B) In female experimental mice, strain and the interaction between strain and diet contribute to variation in body weight. C) In male mice, strain, diet, and their interaction contributed to variation in body weight.

**Table 1.** Results of analyses of body size including all strains with models of the form: Response Variable ∼ Strain * Diet + Covariate.

| Response Variable | Sex | Predictor | Sums of Squares | df | F | P-value |
| --- | --- | --- | --- | --- | --- | --- |
| Body Weight (gm) | M | Strain | 223.757 | 7 | 7.543 | 1.65 x10<sup>-7</sup> |
|  |  | Diet | 51.405 | 1 | 12.130 | 0.001 |
|  |  | Wean Weight | 203.416 | 1 | 48.001 | 2.34 x10<sup>-10</sup> |
|  |  | Strain:Diet | 79.979 | 7 | 2.696 | 0.013 |
|  |  | Residuals | 504.289 | 119 |  |  |
|  | F | Strain | 111.262 | 7 | 8.219 | 3.46 x10<sup>-6</sup> |
|  |  | Diet | 6.205 | 1 | 3.209 | 0.076 |
|  |  | Wean Weight | 157.666 | 1 | 81.533 | 3.11 x10<sup>-15</sup> |
|  |  | Strain:Diet | 42.617 | 7 | 3.148 | 0.004 |
|  |  | Residuals | 235.921 | 122 |  |  |
| Body Length (mm) | M | Strain | 431.397 | 7 | 3.283 | 0.003 |
|  |  | Diet | 5.462 | 1 | 0.291 | 0.59 |
|  |  | Wean Weight | 516.472 | 1 | 27.516 | 6.90 x10<sup>-7</sup> |
|  |  | Strain:Diet | 101.002 | 7 | 0.769 | 0.615 |
|  |  | Residuals | 2252.372 | 120 | NA |  |
|  | F | Strain | 220.139 | 7 | 1.844 | 0.085 |
|  |  | Diet | 52.740 | 1 | 3.093 | 0.081 |
|  |  | Wean Weight | 204.323 | 1 | 11.981 | 0.001 |
|  |  | Strain:Diet | 126.333 | 7 | 1.058 | 0.39 |
|  |  | Residuals | 2097.542 | 123 | NA |  |
| Body Weight/Length | M | Strain | 0.017 | 7 | 6.640 | 1.24 x10<sup>-6</sup> |
|  |  | Diet | 0.005 | 1 | 13.706 | 3.25 x10<sup>-4</sup> |
|  |  | Wean Weight | 0.011 | 1 | 31.405 | 1.38 x10<sup>-7</sup> |
|  |  | Strain:Diet | 0.006 | 7 | 2.368 | 0.027 |
|  |  | Residuals | 0.043 | 119 | NA |  |
|  | F | Strain | 0.011 | 7 | 7.027 | 4.88 x10<sup>-7</sup> |
|  |  | Diet | 0.000 | 1 | 0.680 | 0.411 |
|  |  | Wean Weight | 0.013 | 1 | 58.084 | 6.03 x10<sup>-12</sup> |
|  |  | Strain:Diet | 0.005 | 7 | 3.029 | 0.006 |
|  |  | Residuals | 0.028 | 122 |  |  |
| BMI (kg/m<sup>2</sup>) | M | Strain | 1.254 | 7 | 3.421 | 0.002 |
|  |  | Diet | 0.451 | 1 | 8.603 | 0.004 |
|  |  | Wean Weight | 0.349 | 1 | 6.663 | 0.011 |
|  |  | Strain:Diet | 0.441 | 7 | 1.204 | 0.306 |
|  |  | Residuals | 6.232 | 119 |  |  |
|  | F | Strain | 1.225 | 7 | 4.078 | 4.71 x10<sup>-4</sup> |
|  |  | Diet | 0.002 | 1 | 0.053 | 0.819 |
|  |  | Wean Weight | 0.937 | 1 | 21.837 | 7.713 x10<sup>-6</sup> |
|  |  | Strain:Diet | 0.684 | 7 | 2.276 | 0.033 |
|  |  | Residuals | 5.234 | 122 |  |  |
| Growth Rate | M | Strain | 1.597 | 7 | 9.115 | 5.54 x10<sup>-9</sup> |
|  |  | Diet | 0.347 | 1 | 13.871 | 3.007 x10<sup>-4</sup> |
|  |  | W12 Length | 0.572 | 1 | 22.863 | 5.028x10<sup>-6</sup> |
|  |  | Strain:Diet | 0.433 | 7 | 2.470 | 0.021 |
|  |  | Residuals | 2.978 | 119 |  |  |
|  | F | Strain | 0.739 | 7 | 7.383 | 2.19 x10<sup>-7</sup> |
|  |  | Diet | 0.028 | 1 | 1.991 | 0.16 |
|  |  | W12 Length | 0.071 | 1 | 4.949 | 0.028 |
|  |  | Strain:Diet | 0.273 | 7 | 2.724 | 0.012 |
|  |  | Residuals | 1.745 | 122 |  |  |

In male mice, diet contributed significantly to variation in measures that included body weight (**Table 1**). The same was not true in female mice (**Table 1**). However, in both male and female mice, Tukey’s tests provide evidence that mice on the high-fat diet are significantly bigger (**Table S8**). Importantly, in all tests of measures that incorporate body weight in female mice, the interaction between strain and diet contributed significantly to variation (**Table 1**). Although results were less extreme, that was also true in male mice with the exception of BMI (**Table 1**). Significant contributions to variation from the strain x diet interaction provide evidence that there is variation among the strains in phenotypic response to diet. Thus, there is evidence for genetic variation in plasticity in response to diet amongst these strains. These results are consistent with studies of classical inbred and recombinant inbred strains (e.g., [31,47]) and underscore the importance of using genetically diverse strains in biomedical studies of response to high-fat diet.

There was no evidence of differences among strains in food intake (**Table S9**) consistent with past results [18]. There was also little evidence that diet contributed to differences in food intake, although the strain x diet interaction did contribute to variation in food intake in week 5 in female mice (**Table S9**). Because these strains are new, we also collected testis weights. Strain contributed significantly to variation in testis weight (**Table S9**), but this was largely driven by differences among EDMF males and the other strains (**Table S6**). Despite differences in design and conditions, for strains also reported in Dumont *et al.* ([24]; MANB, SARA, SARB, SARC), average values were broadly similar between the studies.

### Extensive gene expression divergence between strains

To examine gene expression differences in house mice related to metabolic evolution and local adaptation, we analyzed gene expression differences across a subset of strains derived from four localities in the Americas: New York, USA (SARA, SARB), Florida, USA (GAIB), Edmonton, Canada (EDME), Manaus, Brazil (MANB). Results of analyses of body size with this subset of strains were broadly similar to results with the full set (**Fig. S3, Table S10**). To examine the impact of environmental variation on gene expression, liver tissue from each strain was collected from mice on both the high-fat and regular diet. Collections from each biological replicate were completed at 12 weeks post-weaning and mRNA was extracted and sequenced (for details, see Methods).

Strain and sex were the primary drivers of expression variation. Principal component (PC) analysis of gene-wise mRNA abundance separated samples by sex on PC1 and strain on PC2 (27% and 21% of variance, respectively) (**Fig. 2A**). Strains clustered on PC2 consistent with genetic distance [24]. Reflecting their shared geographic origin, the two New York strains (SARA and SARB) clustered closely on PC2. Within sexes and strains, diet treatment was also found to be a major source of variation (**Figs. S4, S5**).

**Figure 2.**
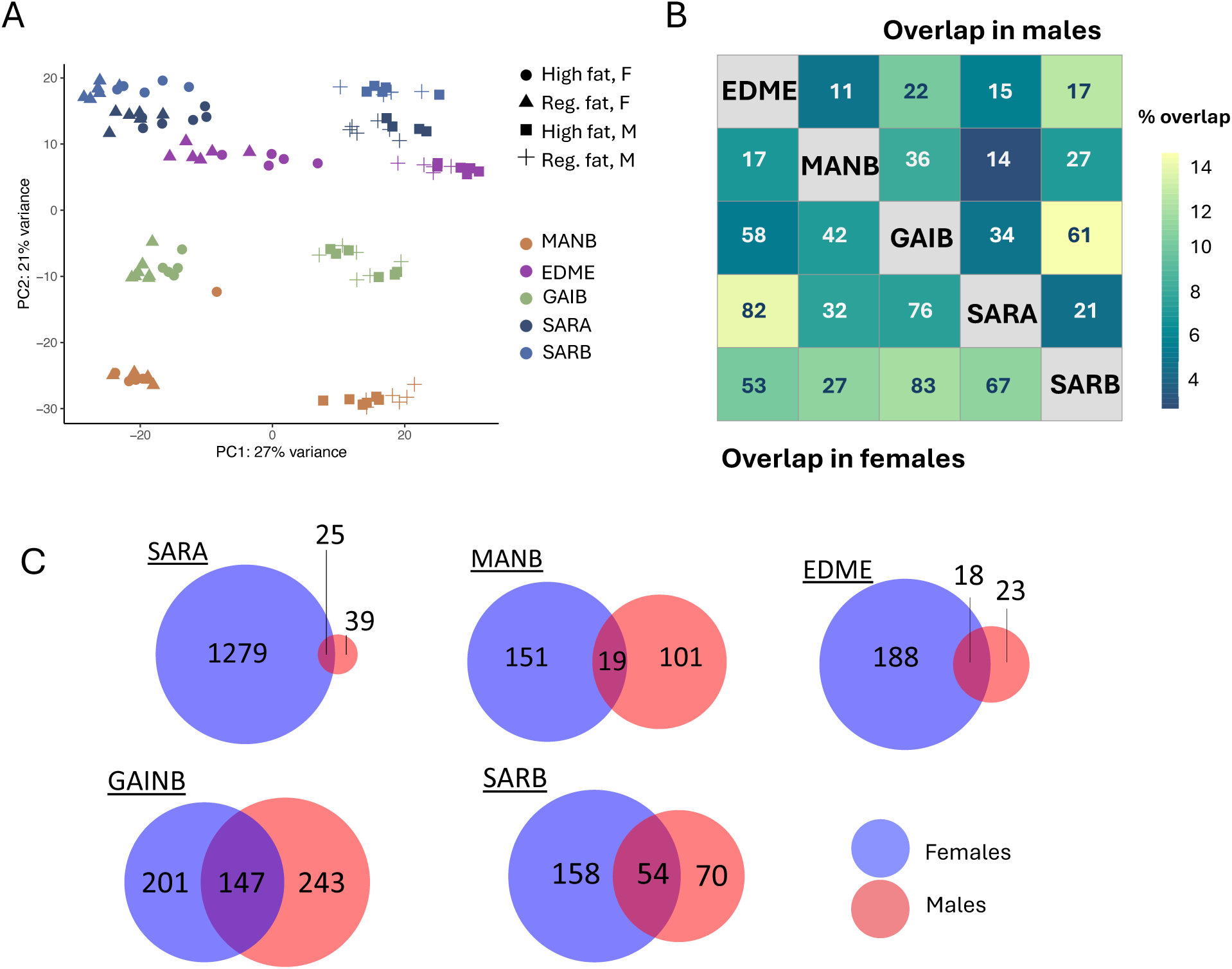
Gene expression differences across populations and diet treatments. **A)** Principal component analysis of gene-wise mRNA abundance. Population is indicated by color, where sex and diet are designated by shape. **B**) Overlap between strains in expression responses to diet. In each square is the number of genes with differential expression between a high-fat and standard diet shared between two strains. Boxes are colored by percent overlap. **C**) Overlap in differentially expressed genes between diet treatments in males and females.

A total of 22,497 genes could be examined for differential expression between strains. Pairwise comparisons revealed extensive gene expression differences between strains. Between 2,389-6,335 of genes showed significant differences in expression between at least two genotypes under either diet (DESeq2 Wald-test) (**Table S11**).

### Gene expression responses to diet are strain-and diet-specific among North and South American mice

Next, we examined gene-by-diet interactions affecting gene expression by comparing mice on a the two diets. To understand whether the transcriptional response to diet is shared or strain-specific, we compared expression between diet treatments for each strain. Across all genotypes, we identified 2,163 genes with a significant transcriptional response to diet, representing approximately 10% of genes surveyed (41-1,203 genes per line/sex; DESeq2 Wald test, FDR<0.05). Genes with changes in expression in response to diet were significantly enriched for GO terms associated with aspects of metabolism (e.g., cholesterol biosynthetic process, triglyceride metabolic process, regulation of fatty acid metabolic process) as well as genes with mutant phenotypes that affected lipid levels (MP:0001547, *q*=3.19x10^-8^), triglyceride levels (MP:0011969, *q*=7.4x10-8), and cholesterol levels (MP:0003947, *q*=0.003).

Surprisingly, diet-treatment effects were largely strain-and sex-specific rather than shared (**Fig. 2B**), indicating genotype-by-environment interactions are pervasive. Over 80% percent of diet-induced changes in gene expression were observed in only one genotype (1,749 genes). Female mice from the SARA line showed the greatest transcriptional response to diet based on the number of genes with differential expression (1,303) and average fold change per gene (|log_2_ (high-fat/regular)|; Kruskal-Wallis, *p*=5.12 x 10^-174^, ad-hoc Wilcoxon signed-rank test, SARA female vs. other genotypes, *P*<2.38x10^-75^). Genes with diet-induced changes in expression specific to the SARA line were enriched for ontogeny terms related to lipid metabolism and gene expression (e.g., lipid metabolic process, *q*=2.2x10^-3^; gene expression, *q=*1.04x10^-5^; rRNA processing, *q*=3.45x10^-2^) relative to other genes with diet-induced expression changes. Between the two strains derived from the New York population (SARA and SARB), we also found transcriptional response to diet to be highly divergent. Only 7.5% and 14% of genes with transcriptional responses to diet were shared between the two lines for females and males, respectively.

Only one gene, *Gstm2,* showed expression changes in response to high-fat diet across all genotypes, though basal expression was significantly different across strains (**Fig. S6**). *GSTM2* expression has previously been shown to protect mice from excess fat accumulation on high-fat diets [48]. Eight genes showed significant responses to diet in males across all lines (*Rnd2, Celf2, Cyp2c29, Cyp2c55, Gstm2, Gstm1, Gstm2-ps1, Gm32342*) and four showed diet-responses in across all lines for females (*Gstm3, Ctse, Mgrn1, Gstm2*). Other genes with diet-responses across multiple genotypes included genes with known roles in metabolism (e.g., *Apoe, Apoa1*, *Irs2, Fabp5, Scp2, Srebf2),* and responsiveness to diet in classical inbred mouse strains (e.g., *Scd1*, *Cyp3a11*) [49,50].

### Stronger transcriptional responses to high-fat diet in female mice

We then examined sex-specific responses to diet affecting gene expression (**Fig. 2C**). Over twice as many genes were found to show diet-based changes in gene expression in females compared to males (1,862 genes and 571 genes, respectively, FDR<0.05). However, 1,107 genes with female-specific diet-responses were observed only in the SARA line. Excluding SARA, a greater number of expression changes were still observed for females, but this difference was more modest (717 vs. 548 genes). The magnitude of expression difference in response to diet per gene was also higher for females than males in all strains but MANB (|log_2_ fold change (High fat/Regular fat)|; Wilcoxon signed-rank test: MANB *P*-value=0.79, other *P*-values=<0.0069). The greatest average expression difference between the sexes was observed for SARA mice, followed by the EDME mice (mean |log2| difference SARA =0.087, EDME=0.02).

Next, we used DESeq2 to identify significant sex-by-diet effects on expression in individual strains. Forty-three genes were found to have significant sex-by-diet interactions. Nearly all of genes with sex-by-diet interactions were strain specific (42/43 genes, FDR<0.05). Genes with significant sex-by-diet interactions in the liver were enriched for metabolic process terms (thyroid hormone metabolic process, *q*=8.75x10^-4^; lipid metabolic process, *q*=1.42x10^-2^; ethanol metabolic process, *q*=2.59x10^-4^).

### Strain-specificity of high-fat diet responses in wild-derived and classical inbred strains

It is clear that morphological response to high-fat diet is highly dependent on genetic background, both from this study and studies of classical inbred and recombinant inbred strains of mice [31,47]. To further examine the strain-specificity of transcriptional response to diet, we re-analyzed liver expression data from Bachmann *et al.* [31] for six classic inbred lines (NZO/HILtJ, C57BL6/J, DBA2/J, A/J, 129S1/SvlmJ, NOD/ShiLtJ) and three wild-derived inbred lines (WSB/EiJ, PWK/PhJ, CAST/EiJ) subjected to either a high-fat or standard diet (see Methods). In this dataset, between 33-2,447 genes were differentially expressed between diet treatments for each genotype (0.02-14% of genes surveyed) (See Methods). As in our dataset, more differentially expressed genes were identified in female mice (5,864 vs. 3,026 genes) [31].

Approximately 54% of genes with diet-based differences in expression in our dataset were also identified in at least one other classical or wild-derived inbred strain [31] (1,171 genes). This degree of overlap was observed despite differences in time spent on a high-fat diet and age across experiments (see Methods). Comparing across all strains, 8,359 genes showed significant differences in expression associated with diet. Importantly, though, approximately 58% of these were observed in only one genotype. Overall, both classical and wild-derived inbred lines show high levels of strain-specificity in their transcriptional response to diet (**Figs. S7, S8, S9**). These results emphasize the importance of GxE interactions in both evolutionary and biomedical contexts.

### Gene co-expression analysis identifies genes associated with variation in metabolic traits

To investigate the associations between gene expression and metabolic traits, we performed a weighted gene co-expression network analysis (WGCNA) [51]. WGCNA was used to identify groups of genes with highly correlated expression, called co-expression modules, for each diet treatment and sex separately. Co-expression modules were then tested for correlations with metabolic phenotypes (body weight, body length, BMI, growth rate, and food intake for weeks 1 and 9). Metabolic phenotypes, except for measures of food intake, were non-independent and correlated across individuals (**Table S12**). We identified several modules associated with trait variation in each group (**Table S13**). Between 9-17 co-expression modules were associated with body mass in each comparison, the majority of which were also significantly associated with other metabolic traits. In contrast, few co-expression modules were associated with food intake measures (2-4 modules per diet and genotype combination) and these modules were typically not significantly associated with other metabolic phenotypes.

To identify common modules across diet treatments, we performed a consensus network analysis (full dataset available in Supplemental material online). In a consensus network analysis, co-expression modules are identified across treatments (63 in males, 32 in females). We saw relatively high network preservation across diets (Density value (D)=0.92 in females, 0.83 in males). In the consensus network, we identified co-expression modules associated with weight (7 and 14 modules in females and males, respectively), BMI (2 and 3), body length (4 and 15), growth (5 and 11), and food intake (1 and 0) (**Figs. S10, S11**). To identify modules with different associations with metabolic traits in high-fat and standard diets, we compared the preservation between module and trait associations (i.e., weight, BMI) across these sets. Overall, the relationship between traits and modules were highly preserved across diets (**Figs. S12, S13**). This result suggests that, while at finer resolution transcriptional response to diet is highly strain specific, at the level of pathways, associations between phenotype and expression are largely shared. However, we did identify module-trait relationships with lower preservation, where a module was correlated with trait variation under only one diet. For example, in males the “yellowgreen” module was associated with body mass only under a high-fat diet (high-fat, corr=0.69, *p*=4 x 10^-9^; standard, corr=-0.16, *p*=0.4, preservation=0.57). Genes in this module with strong associations with body weight under the high-fat diet include *Insulin induced gene 2 (Insig2)*. Genetic variation at this gene has been associated with metabolic traits in humans [52–54]. In females, the “steelblue” module was associated with body mass variation only under a high-fat diet (high-fat, corr=0.63, *p*=2 x 10^-4^; standard, corr=0.21, *p*=0.3, preservation=0.79).

Genes in this module were enriched for ontology terms related to cell cycle (e.g., cell cycle, *q*=1.29x10^-41^, cell division, *q*=7.66 x 10^-38^). Therefore, these modules may contain genes involved in phenotypic responses specific to high-fat diet.

### Gene regulatory divergence between mice from divergent climates

Because we wanted to study gene expression regulatory divergence, we also crossed strains from locations with different climates (GAIB x SARA, SARB X MANB; **Fig. 3A**) and subjected progeny to the same experimental manipulation described above. Body size measures were collected (**Table S3, Figs. S14, S15**) and liver tissue was collected for mRNA sequencing.

**Figure 3.**
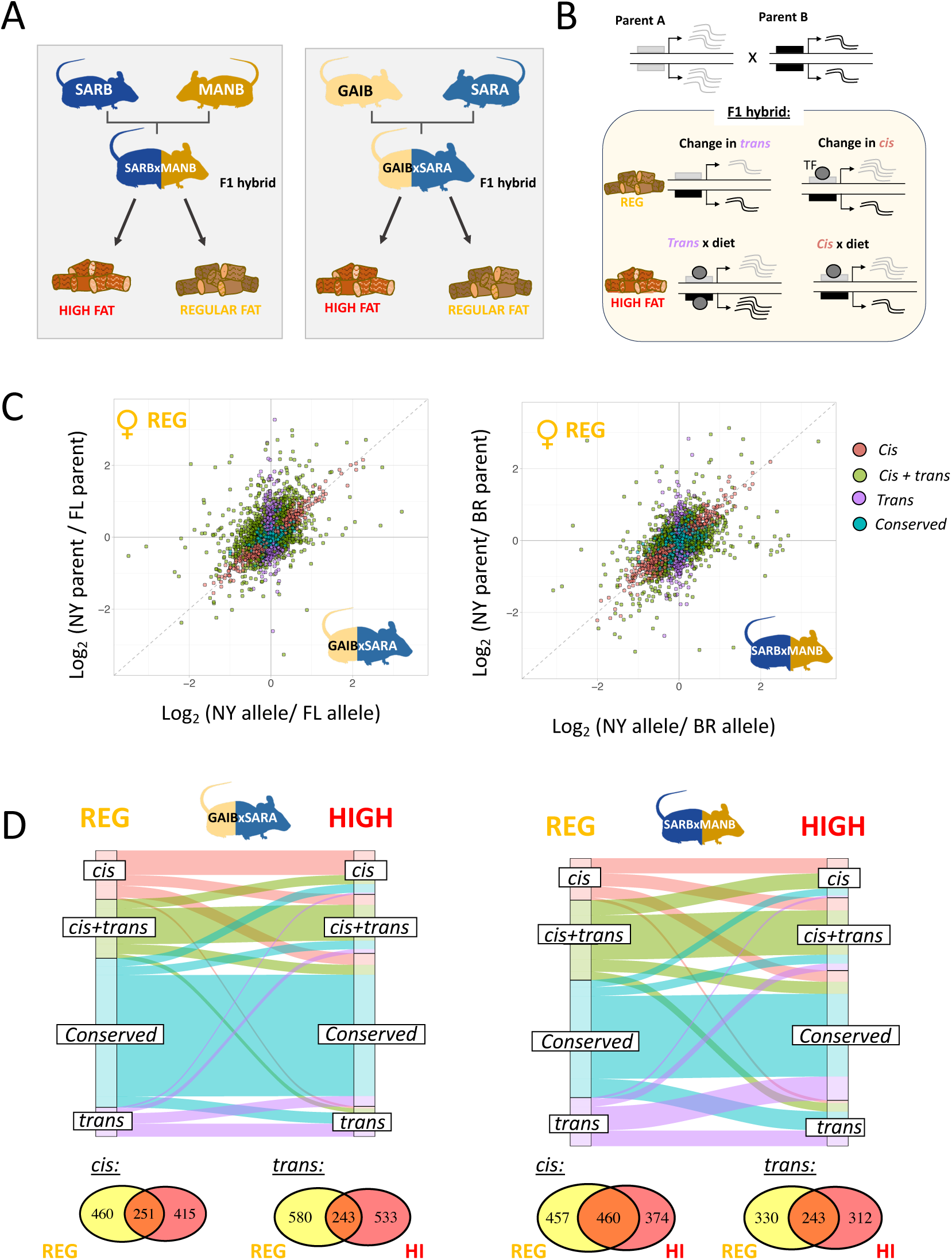
Gene expression regulatory divergence across strains and diet treatments. **A**) Mice from different localities were crossed to produce F1 hybrids. Parental lines and F1 hybrids were raised on either a regular or high-fat diet. Livers were collected for RNA-seq for at least four replicates per group (population, sex, diet). **B**) A schematic showing how regulatory differences can be inferred via allele-specific expression. **C**) Gene expression regulatory divergence between New York (SARA) and Florida (GAIB) strains (left) and New York (SARB) and Brazil (MANB) (right) was characterized through comparisons of allele-specific expression in hybrids and parental gene expression for female mice. Each point indicates a gene colored by their regulatory classification. **D**) Comparison of gene regulatory divergence between female mice on high-fat(“HIGH”) and regular (“REG”) diets. Comparison across diets revealed extensive environmental effects on gene regulation. Venn diagrams show the number of genes with *cis* vs. *trans* effects across diet treatments.

Expression measurements in F1 hybrids can be used to discriminate between local regulatory changes (*cis*-divergence) vs. distant regulatory changes (*trans-* divergence). As F1 hybrids inherit a copy of each chromosome from both parents, alleles from both parents are present in the same cellular environment. Consequently, differences in expression between the two parental alleles in a hybrid (i.e., allele-specific expression; ASE) is indicative of a *cis-*regulatory change. In contrast, when differences in expression are observed between parents but not in F1 hybrids, we can infer that expression differences are due to *trans-*acting changes (**Fig. 3B**) [55].

Gene regulatory divergence was examined in 6,951 and 5,716 genes between SARB and MANB and GAIB and SARA, respectively (**Fig 3C**) (see Methods). Between SARB and MANB, *cis-* changes were found to be predominant over *trans-* changes, consistent with previous intraspecific studies of house mice (**Table S14**). In contrast, between GAIB and SARA, *trans-* changes were typically found to be predominant **(Table S15**). As *cis-* regulatory changes have been found to accumulate with genetic distance in other systems, this difference may reflect greater divergence between the New York, USA and Manaus, Brazil populations compared to New York, USA and Florida, USA populations [56]. Effect sizes of *cis-*changes were also found to be smaller on average between SARA and GAIB than SARB and MANB, again potentially reflecting lower genetic divergence between these populations (Wilcoxon signed-rank tests, *P-* values<0.01).

### Gene regulatory divergence between populations is diet-dependent

The environment can induce changes in gene expression via *cis-* or *trans-*acting mechanisms (**Fig. 3B**). To understand the relationship between environment and gene regulatory divergence, we compared our results between individuals fed a regular vs. high-fat diet. We found that gene regulatory divergence was often dependent on diet. Between 32-45% of genes showed a different regulatory categorization between diet treatments for each comparison (**Fig. 3D**). This suggests that gene regulatory divergence between strains is highly context specific. The greatest differences in regulatory categorization were observed between females from the GAIB and SARA cross, likely reflecting the unique response of SARA females to diet (see above, **Fig. 2C**). A greater proportion of genes showed evidence of regulatory divergence in *cis* and/or *trans* on a regular diet in most comparisons (Fisher’s exact tests, SARB x MANB females, *P*=0.25, all other comparisons *P*<0.0003), suggesting a greater conservation of regulatory architecture between strains on a high-fat diet.

Overall, we found that *cis-* regulatory divergence between strains was more robust to diet, where *trans-*divergence was more responsive to diet differences. Gene expression differences ascribed to *cis-* changes were more stable across diet treatments than *trans-* changes in most comparisons (Fisher’s exact tests, GAIB x SARA Females *P*=0.07, all other crosses *P*<0.0001). Effect size differences also reflected this, with diet having a smaller effect on *cis-* compared to *trans-* divergence (Wilcoxon signed-rank tests, *P-*values<7.27 x10^-113^). The robustness of *cis*-acting changes compared to *trans-*acting changes to environmental variation is consistent with studies in other systems [25,57–59]. Overall, evidence suggests that *trans* regulation may play a more important role in environmentally induced changes in gene expression.

### Evidence for polygenic selection on *cis*-regulatory divergence between mice from divergent climates

Previous work in this system suggests *cis-*regulatory changes in particular may play an important role in environmental adaptation [18,25–27]. Gene expression adaptation via *cis*-regulatory mutations can be highly polygenic, involving many independent changes at functionally related genes [60–62]. To test for evidence of selection on *cis-*regulatory changes between mice from divergent climates, we applied a gene-set test of selection based on a sign test framework [61,62]. For a given pathway or biological function, we expect *cis*-regulatory changes to be unbiased with respect to their directionality (e.g., an equal number of *cis*-changes upregulate SARB and MANB alleles). If, however, *cis-*regulatory changes act in the same direction more than expected, this can indicate lineage-specific selection on the *cis*-regulation of this gene set (**Fig. 4A**). Applying the sign test to GO gene sets in each cross, we identified multiple gene sets with evidence for biased directionally (full list in **Table S16**) (see Methods).

**Figure 4.**
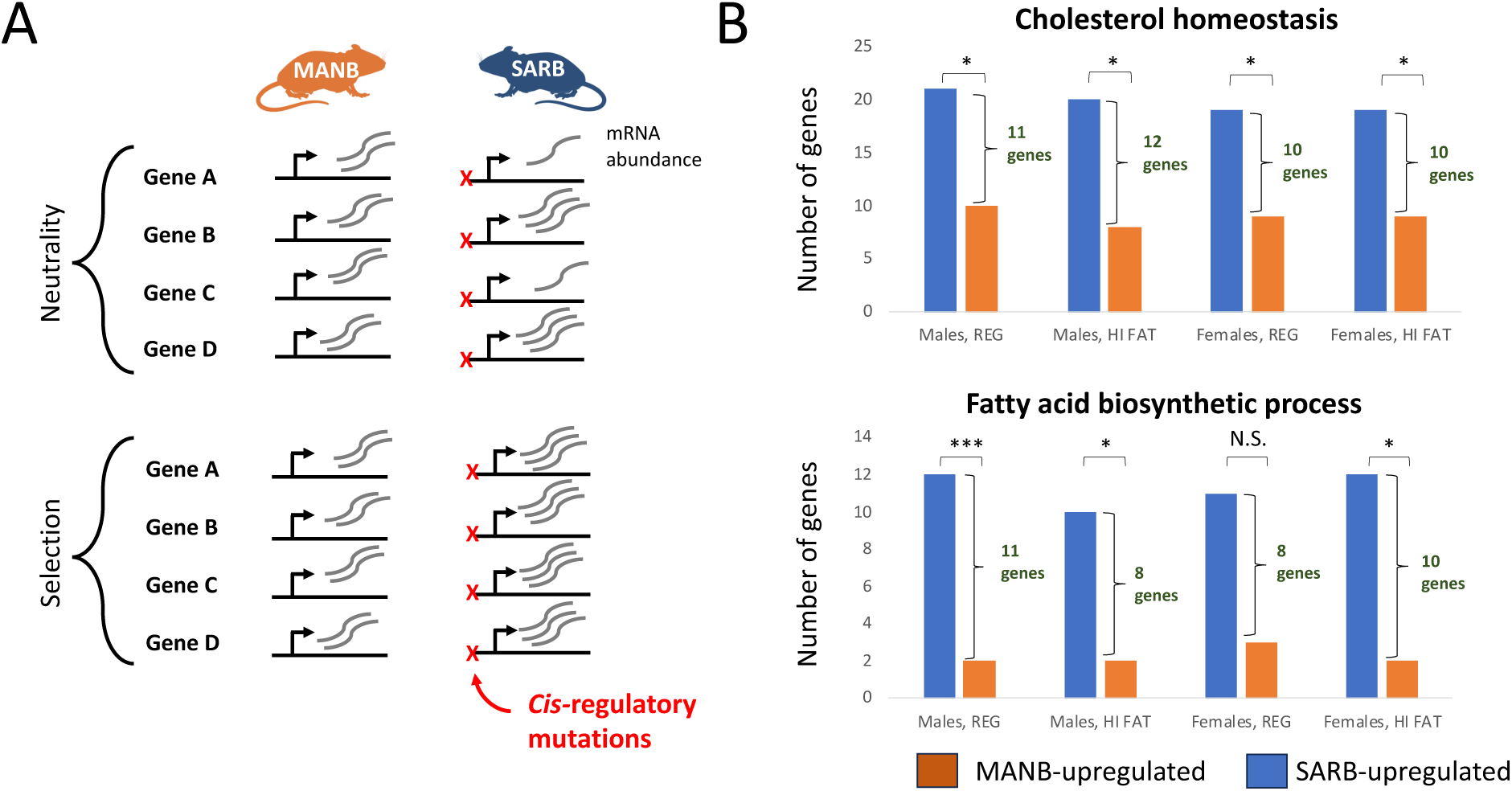
Evidence for polygenic *cis*-regulatory evolution. **A**) Testing for polygenic *cis-* regulatory evolution (adapted from [62]). Examples of expression (squiggly lines) of four functionally related genes are shown for MANB and SARB. An “X” indicates an independent *cis-* regulatory mutation that alters expression. Under neutrality, we expect an equal number of *cis-* regulatory changes affecting expression to increase and to decrease expression as shown in the top panel. However, if in comparing MANB and SARA, we see that all *cis-*regulatory mutations increase expression of the SARB allele (as in the bottom panel), this would be consistent with lineage-specific selection for altered expression of this entire gene set. B) Effect directions of ASE for genes related to cholesterol homeostasis (top) and fatty acid biosynthetic process (bottom). We observed a significant bias of upregulation of SARB-alleles (blue bars) vs. MANB-alleles (orange bars) for genes associated with these processes. \*\*\**P*<0.001, \*\*\**P*<0.01, N.S. = Not significant.

In the SARB x MANB cross, multiple GO gene sets related to metabolism (Fatty acid biosynthetic process, Benjamini-Hochberg adjusted *P* (*q*)=0.0058; Cholesterol homeostasis, *q*=0.079; Retinol metabolic process, *q=*0.079) showed significantly biased upregulation of New York alleles compared to Manaus alleles (**Fig. 4B**, **Table S16** and Supplemental online data). Genes associated with metabolism ontogeny terms include important regulators of body weight and fat accumulation (e.g., *Fads2, Acsm1, Acsm3, Scd1*) [63–65]. Polygenic selection on *cis*-regulatory variation related to metabolism is consistent with phenotypic differences in body mass and blood chemistry observed between mice from divergent climates [18,20]. Other gene sets with evidence for biased directionality included terms related to immunity (e.g., Adaptive immune response, *q*=0.0058; Antibacterial humoral response, *q*=0.008) and cellular processes (e.g., regulation of apoptotic process, *q*=6.21x10^-5^). In the GAIB x SARA cross, only one GO term (Response to ethanol, *q*=0.0046) was significant after correction for multiple testing.

### *Cis*-by-diet interactions are common and related to metabolism

Next, we formally tested for context-specific *cis*-regulation associated with diet. To identify specific genes with evidence for *cis*-by-diet effects on expression (**Fig. 3B**), we compared ASE ratios in hybrids across diet treatments (i.e., GAIB/SARA allele in high-fat vs. GAIB/SARA allele in regular and MANB/SARB allele in high-fat vs. MANB/SARB allele in regular). We identified 425 genes with a *cis*-by-diet interaction (See Methods, FDR<0.05) (**Fig. S16**). The majority of *cis*-by-diet interactions were observed in only one cross or sex (∼81%), with only three genes showing *cis-*by-diet effects in all comparisons (*Serpinc1*, *Fmo5*, *B2m*). Consistent with a potential role for these changes in modulating responses to diet between strains, *cis*-by-diet interactions were enriched for metabolic GO terms (e.g., fatty acid metabolic process, cellular response to cholesterol, regulation of bile acid secretion; FDR<0.05) and phenotypes (e.g., abnormal triglyceride level, abnormal phospholipid level, abnormal food intake; FDR<0.05) compared to all genes with ASE.

On average, *cis-*by-diet interactions were of small or modest effect that resulted in a change in the magnitude of ASE between lines (**Fig. S16**). However, we also identified cases where *cis*-changes were specific to one diet (122 genes). We consider these cases of diet-induced allele-specific expression. Many of these genes have previously been linked to diet-induced metabolic phenotypes in either humans or mice (Supplemental online data).

Genes with *cis*-by-diet interactions were also identified in gene co-expression modules associated with metabolic phenotypes. These genes are promising candidates for mediating strain-specific diet responses (Supplemental online data). For example, protein sterol carrier protein-2 (*Scp2*) is a hub gene in the module with the strongest positive association with body weight in males on a high-fat diet (green module, corr=0.83). *Scp2* was found to have a *cis*-by-diet interaction in the SARB x MANB cross and is significantly correlated with BMI and body weight variation in males in our data (standard diet, corr=0.6, *p*=0.0006; high-fat diet, corr=0.83, *p*=8.73 x 10^-8^). *Scp2* is involved in regulating lipid metabolism in the liver[66].

### Overlap between signatures of natural selection and gene expression divergence

We next investigated overlap between our results and genomic signatures of environmental adaptation in North and South American house mice. To examine genomic signatures of environmental adaptation, we utilized previously generated population genomic data for individuals collected from populations in North and South America along a latitudinal transect (134 individuals) [18,20]. Variants associated with latitude were identified using a latent factor mixed model (LFMM) [67] accounting for population structure (as previously described [18,20]). Genomic outliers for associations with latitude for North American populations and South American populations were retained at a z-score >|3| for overlap analysis with the expression data (see Methods).

Differentially expressed genes between the Manaus (MANB) and New York (SARA, SARB) strains overlapped outliers for genome-environment associations in North and South America (223 and 2,502 genes, respectively). Of these, 549 genes showed evidence for *cis*-regulatory divergence between the New York and Brazil strains based on allele-specific expression. These genes are particularly interesting as candidates for local adaptation, as evidence for *cis*-regulatory divergence suggests that there is local genetic variation affecting gene expression.

ASE genes containing genomic outliers were enriched for ontogeny terms related to metabolism (e.g., long-chain fatty acid metabolic process, *q*=7.49x-10^-3^; unsaturated fatty acid metabolic process, *q*=4.51x10^-2^) and gene expression regulation (e.g., regulation of transcription by RNA polymerase II, *q*=5.36x10^-6^) over the background of genes that could be tested for ASE. Included in this set are genes identified in our ASE sign test (see above, Supplemental online data). For example, several genes associated with the ontogeny term for “fatty acid biosynthetic process” also overlapped outlier loci (e.g., *Acsm1, Elovl5, Elovl2*) as did genes associated with “cholesterol homeostasis” (e.g., *Lipc, Ttc39b, Nr1h4*). In comparisons between New York and Florida, 246 genes with evidence for differential expression overlapped genomic outlier regions in North America, of which 59 were identified as having a *cis-*component underlying regulatory divergence. We found no enrichment for this set.

Highlighting the importance of examining gene expression under multiple environmental contexts, 110 genes overlapping outlier loci also showed cis-*by*-diet interactions. This set included genes with known roles in gene-by-diet effects (**Table 2**) and was also enriched for metabolic GO terms (e.g., fatty acid metabolic process, 6.29 x 10^-3^; lipid metabolic process 2.30 x 10^-2^) relative to other ASE genes overlapping genomic outliers (Supplemental online data).

**Table 2.**
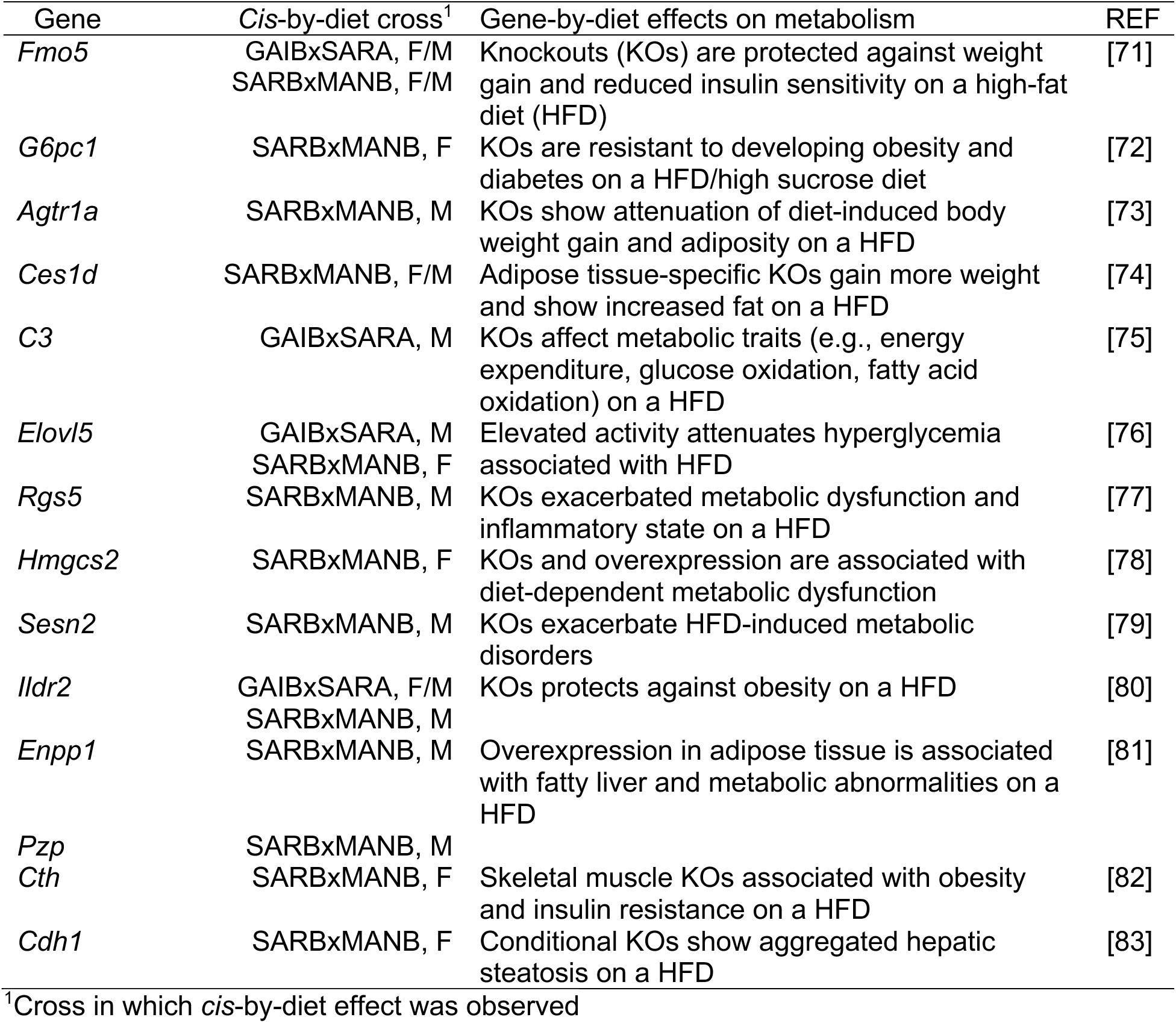
Genes with *cis*-by-diet effects overlapping outlier loci with diet-dependent effects on metabolism.

Differential gene expression and ASE can help make connections between candidates from genome scans and adaptive phenotypic variation (e.g., [18,25,26,68–70]). Incorporating environmental effects further ties candidates to phenotype by identifying induced differences in gene expression and *cis*-by-environment interactions.

### Conclusions

The evolution of complex traits is inextricably tied to the environment. In this study, we leveraged a new set of wild-derived inbred mouse strains from divergent climates, using an experimental manipulation of diet to investigate the role of the environment and GxE interactions in shaping adaptive variation in complex traits. We found that there were significant effects of strain, diet, and strain-by-diet interactions on body size and gene expression. In fact, transcriptional response to diet was largely strain-specific, making GxE interactions the rule rather than the exception.

Using crosses between strains from differing climates, we were able to further strengthen ties between gene regulation, the environment, and adaptation. We identified both *cis-* and *trans-* acting changes that contributed to gene expression differences between strains from tropical and temperate regions of the Americas. Importantly, we found that gene regulatory divergence is highly dependent on environmental context, particularly for *trans*-acting changes. Higher sensitivity of *trans*-divergence to diet points to a greater role for *trans* regulation in gene expression plasticity. In contrast, associations between metabolic phenotypes and gene co-expression modules were largely robust to changes in diet, highlighting the exceptions as modules containing candidate genes for phenotypic response to diet. We were further able to connect local variants to plasticity in gene expression and phenotype via identification of genes with *cis*-by-diet interactions that were found in co-expression modules associated with variation in body mass related phenotypes and as a group enriched for metabolic GO terms. Notably, ASE sign tests provide evidence of selection on *cis-*regulatory changes related to metabolism, consistent with the differences in body mass and blood chemistry among these populations [18,24]. Overlap among outliers from selection scans and differentially expressed genes also helps connect candidate genes to adaptive variation in phenotype, in particular those with evidence of *cis*-regulatory divergence and functional ties to metabolism. Outlier loci with cis-*by*-diet interactions are strong candidates for adaptive plasticity.

More generally, our results underscore the value of incorporating environmental variation in evolutionary and biomedical studies of complex traits. Moreover, they add to a growing body of evidence that genetic variation is crucial to studies of response to high-fat diet, a trait of considerable biomedical significance (e.g., [31,47]). Most of the strains included in this study are now available commercially making them potentially useful tools for the study of diet-induced obesity [24]. Sex specific transcriptional responses to diet also point to the importance of explicitly considering sex both in experimental design and in investigating underlying biological processes. Overall, this study represents a significant step forward in our understanding of adaptive body size variation in this system and demonstrates the importance of environmental variation and GxE interactions in disentangling the processes that underlie adaptive variation in complex traits.

## Methods

### Experimental Design

To evaluate variation in response to a high-fat diet, we selected eight newly derived strains of *M. m. domesticus*. Four were derived from populations in the United States: TUCC (Tucson, AZ), GAIB (Gainesville, FL) and SARA, SARB, and SARC (Sarasota Springs, NY). The remainder were from Canada, EDME and EDMF (Edmonton, Alberta) and Brazil, MANB (Manaus, Amazonas). The strains were generated via brother-sister pairing [18,24] for at least 12 generations. Importantly, these mice descend directly from wild-caught progenitors and were not rederived, a process which would be expected to alter their gut microbiome. However, several of these strains have now been rederived and are available commercially [24]. Mice were kept in common laboratory conditions at approximately 22℃ with a 14/10 light/dark cycle throughout the experiment. After weaning at ∼24 days, mice were housed singly with enrichment and provided with water *ad libitum* as well as one of two diets: high-fat (*TestDiet* BLUE 58Y1, 34.9% fat) or a breeder chow (*LabDiet* Extruded Picolab select mouse diet SVSM 50 IF/9F, 8% fat). At weaning and every week thereafter, animals were weighed (SPX222 Scout Analytical Balance, Ohaus, Parsippany, NJ, USA). Body length and tail length were measured via ruler at 1, 5, and 9 weeks. In addition, at those time points, ∼120 grams of food was weighed, placed in a clean cage, and collected and re-weighed 24 hrs later (SPX222 Scout Analytical Balance, Ohaus). From these data, the amount of food consumed per hour was calculated. In some cases, food was weighed slightly earlier or later and, in those cases, the rate calculations were corrected for the difference. In addition, many mice ground food excessively, resulting in food in the cage that was clearly not consumed. In these cases, food intake data were not included in the analysis. After 12 weeks, animals were euthanized. Body weight, body length, tail length, hindlimb length, and ear length were measured (SPX123 Scout Analytical Balance, Ohaus) and liver tissue was immediately collected for gene expression analyses. Livers were stored in RNAlater at 4℃ for ∼24 hours and then transferred to a -80℃ freezer. All work was performed with approval from the Monmouth University Institutional Animal Care and Use Committee.

RNA was extracted from frozen liver tissues using the Qiagen RNeasy+ Mini kits with tissue disruption via bead beating (Beadruptor 12, Omni International, Kennesaw, GA, USA). RNA samples were quantified (Qubit 2.0 Fluorometer; ThermoFisher Scientific, Waltham, MA, USA) and integrity was verified (4200 TapeStation; Agilent Technologies, Palo Alto, CA, USA). RNA quality was high (RIN_average_=9.90, sd=0.25) and strand-specific libraries were prepared (Azenta, Inc., South Plainfield, NJ) using NEBNext Ultra II RNA Library prep kits (New England Biolabs, Ipswich, MA, USA). Sequencing libraries were validated (Agilent TapeStation) and quantified via Qubit as well as by quantitative PCR (KAPA Biosystems, Wilmington, MA, USA). For the initial run, the sequencing libraries were multiplexed and clustered onto 11 flow cells. After clustering, flow cells were loaded onto the Illumina HiSeq instrument and sequenced using a 2x150bp Paired End (PE) configuration. Raw sequence data (.bcl files) were converted into fastq files and de-multiplexed using Illumina bcl2fastq 2.20 software. One mismatch was allowed for index sequence identification. To augment these data, two additional lanes were sequenced using the same approach on the NovaSeq platform (2x150bp PE). Reads sequenced per sample are available in **Table S17** (median=62,289,663 paired end reads).

### Analysis of morphological data

To test whether aspects of body size varied among strains, we used linear models including relevant factors, interactions, and covariates (Tables 1, S4-S10). The models were implemented in R via the aov function and evaluated via the Anova function choosing the Type III option [84,85]. Tukey’s tests for each factor were used to further clarify differences. Tests were implemented in R using the TukeyHSD function [86,87].

### Read mapping and parental expression analysis

Reads to the *Mus musculus* reference genome (GRCm39) using STAR (v 2.7.11.a) [88] with the Ensembl GRCm39 annotation. Reads overlapping exons were counted for gene-wise quantification using HTSeq [89]. Raw read count data was transformed using variance stabilizing transformation to assess transcriptome-wide expression patterns via PCA. DESeq2 was used to assess differential expression by fitting a generalized linear model following a negative binomial distribution [90]. Gene expression was compared between lines and between diet treatments using a model that included population of origin, sex, and diet (Wald Test). A Benjamini–Hochberg multiple test correction was used on *P*-values. We considered genes with an FDR < 0.05 to be differentially expressed between comparisons.

### Identifying variants for allele-specific expression

To quantify ASE, we mapped reads from each F1 individual to the mouse reference genome with STAR. We utilized the Genome Analysis ToolKit (GATK) [91] to identify variants for measures of ASE. First, duplicates were marked with the Picard tool MarkDuplicates. Read groups were added with AddOrReplaceReadGroups. SplitNCigarReads was then used to split reads that contain Ns in their cigar string (e.g., spanning splice events). HaplotypeCaller and GenotypeGVCFs were utilized for joint genotyping across parental and F1 samples. SNP calls were filtered with VariantFiltration (QD < 2.0; QUAL < 30.0; FS > 200; ReadPosRankSum < -20.0). Variants were included for downstream analysis if all genotyped F1s were called as heterozygous and parental lines were assigned alternative homozygous calls. We used these calls to create a high-quality list of informative variants for ASE.

To identify reads overlapping informative sites, we mapped F1 reads to the mouse genome using the WASP implementation in STAR to mitigate mapping bias associated with the reference allele[92,93]. WASP eliminates reads with potential for mapping bias by flagging reference biased reads. First, reads containing SNPs are identified. Then, WASP simulates reads with alternative alleles at that locus and remaps reads to the reference. Reads that do not map to the same location are flagged[92]. Reads overlapping informative variant calls that passed WASP filtering were then retained to estimate ASE. HTSeq-count[89] was used to count reads associated with each gene for each parental allele separately.

To identify genes with evidence for allele-specific expression (ASE), reads that mapped preferentially to one parental allele in hybrids were compared with DESeq2[25,68,94]. Reads were fit to a model with allele, sample, and sex (Wald test, FDR<0.05). For hybrid samples, DESeq2 library size factor normalization was disabled as read counts came from the same sequencing library (setting: SizeFactors = 1). Previously published code is available on Github (github.com/malballinger/BallingerMack_PNAS_2023) for these analyses.

### Characterizing gene regulatory divergence

Pure strains (“parental”) and F1 hybrid expression were compared to identify *cis* and *trans* divergence at each gene. Genes were partitioned into different regulatory categories by comparing allelic expression in F1s, expression differences in the parentals, and a comparison between allelic and parental ratios[95]. To equalize power across comparisons, we randomly dropped individuals to maintain an equal number of parental and allelic samples in each comparison. Parental reads were then downsampled to match that of hybrid libraries and all libraries were downsampled to equalize power across replicates [25,56,96]. Parental and allele-specific hybrid counts for each replicate were summed and binomial exact tests were used to identify *cis-* and *trans-* divergence, as in previous studies (see below for method comparison)[25,56,96,97]. Fisher’s exact tests were used to compare allelic and parental ratios. The resulting *P-*values from binomial and Fisher’s exact tests were corrected following the Benjamini–Hochberg method and considered significant at an FDR<0.05. Genes were then sorted into regulatory categories using previously described criteria [25,56,95,96,98]. In brief, genes were sorted as follows:

*Cis* only: (1) a significant difference in expression between populations, (2) a significant difference between alleles in hybrids, (3) no significant difference between parental and allelic ratios.

*Cis+trans:* (1) a significant difference in expression between populations, (2) a significant difference between alleles in hybrids, (3) a significant difference between parental and allelic ratios.

*Trans* only: (1) a significant difference in expression between populations, (2) no significant difference between alleles in hybrids, (3) a significant difference between parental and allelic ratios.

Conserved or Ambiguous: All other patterns.

The downsampling and pooling of replicates was chosen as the approach to categorize genes based on *cis-* and *trans-* divergence due to comparisons with unequal read depth and replication[56]. As in previous studies [25], we saw congruence between the use of binomial tests and DESeq2 for identifying *cis*-regulatory divergence via ASE (DESeq2 Wald-test, see above). *P-*values from DESeq2 and binomial tests were found to be highly correlated (Spearman’s rho>0.79, *p-values*<2.2 x10^-16^ for all pairwise comparisons).

### Comparing liver expression responses to diet between classical and wild-derived inbred strains

Previously published RNA-seq data from Bachman et al. [31] for 9 lines was downloaded from the GEO expression database under the accession GEO:GSE182668. Reads were mapped to the *M. musculus* reference genome (GRCm39) and reads overlapping exons were counted as described above. Count data was then analyzed in DESeq2 using a model incorporating sex, strain, and diet, as described above. We note that differences in experimental approach and other potential batch or technical effects (e.g., sample preparation, sequencing chemistry) likely affect overlap between genes identified between our studies. Experimental differences between the two studies include the following: (1) time and age at which mice are put on a high-fat diet (8 to 21 weeks in Bachmann et al. [31]), (2) age at which livers are collected (>21 weeks vs. ∼15 weeks).

### Weighted Gene Co-expression analysis

To identify co-expression gene sets and associate expression variation with phenotypic variation across strains, we used WGCNA. This approach has previously been used to associate gene expression variation with variation in complex traits and adaptive phenotypes (e.g., [99–101]). We carried out a weighted gene co-expression network analysis on normalized expression data following WGCNA protocols [51,102]. We filtered parental data for genes with low expression across samples (>20 reads per sample in at least four samples), resulting in 15,622 genes for analysis.

We constructed a gene co-expression network, represented by an adjacency matrix, which denotes co-expression similarity between pairs of genes among different individuals. Modules were identified using unsupervised clustering. Dissimilarity between clusters is measured based on the topological overlap and defined by cutting branches off the dendrogram [51,103]. Soft-thresholding power was chosen based on the pickSoftThreshold function in WGCNA to achieve an approximately scale free topology. Expression modules were inferred for standard and high-fat treatment and male and female mice separately, and then consensus modules were created to identify co-expression patterns shared across groups. We chose a minimum module size of 30 genes. To associate modules with traits of interest, we then tested for correlations between eigengenes (the first principal component of a module) and each trait (Pearson correlation). In order to explore the preservation of module trait relationships in standard and high-fat diet treatments, we analyzed the differential eigengene network (Figs. S12, S13). Plots of differential analysis were created with the command *plotEigengeneNetworks*. The density (*D)* of the eigengene network is defined as the average scaled connectivity. Larger values of D (closer to 1) are indicative of stronger correlation preservation between all pairs of eigengenes across the high-fat and standard networks. Gene module membership and trait associations from our consensus WGCNA analysis can be found on FigShare (https://figshare.com/s/721a12ffbe1bfa3b921c).

### ASE sign test

Gene Ontology (GO) categories were obtained through Ensembl annotations for mice (GRCm39). We restricted our analysis to GO Biological Process terms. For each cohort separately (e.g., cross (GAIBxSARA or SARBxMANB), diet-treatment (high-fat, standard) and sex (males, females)), genes with significant evidence for ASE were divided based on which allele was upregulated (SARA/B or MANB/GAIB). GO Biological Process terms with fewer than 10 members in a cohort were excluded from the analysis. A Fisher’s exact test was used to identify lineage-specific bias on each set [62,68,96].

*P*-values for each gene set in a cohort (sex, diet) were combined using Fisher’s method to obtain a single *p*-value for each gene set in a cross (GAIBxSARA or SARBxMANB), as previously described [62,68]. FDR was estimated in two ways, via (1) the Benjamini-Hochberg Procedure implemented in R with *p.adjust*, and (2) gene category assignments permutations [62,68,104]. For each cohort, gene-GO category assignments were randomly shuffled and the test was repeated 1000 times. *P*-values were determined by asking how often a result of equal or greater significance was observed in the permuted set (equivalent to a GO category-specific FDR [104]). Gene sets for which significant biased directionality in both tests are reported in **Table S16**.

### Overlaps with selection scans

Latent Factor Mixed Model (LFMM)[67] software was used to identify associations between genetic variants and latitude as previously described [18,20]. In brief, LFMM was run separately for individuals from east coast populations in the United States [18] and for South American populations(burn-in = 100,000 and 500,000 iterations) [20] along two latitudinal transects. K values of 2, and 3 were used for North America and South America, respectively. For each run, the genomic inflation factor was estimated (λ) and *p*-values were corrected for false-discovery rate. We use outliers at a |z-score| threshold of >3 to examine overlap with expression data.

### Enrichment analyses

Gene Ontology (GO) and pathway enrichment were performed with PANTHER [105]. Phenotype enrichment analyses were performed with ModPhea [106].

## Data Availability

All raw sequence data will be available online through NCBI Bioproject (PRJNA1164275). Supplemental online datasets are available through FigShare (https://figshare.com/s/721a12ffbe1bfa3b921c).

## Acknowledgements

We thank Michael Nachman for graciously sharing the new wild-derived inbred strains used in this study. We thank Yocelyn Gutiérrez-Guerrero for providing data for our analysis of overlap with selection scans. We thank Katherine Banfitch, Erin Oscar, Julia Panebianco, Caroline Reverendo, and Summer Shaheed for husbandry assistance. Research was supported by internal funding from Monmouth University and Drexel University. MPR is supported by NSF Division of Environmental Biology Award #2332998. NL, TL, SV, and LC were supported by the undergraduate summer research program at Monmouth University. KLM is funded by the National Institute of General Medical Sciences of the National Institutes of Health under Award Number P30 GM145499 and NIH R35GM154966. This work used Expanse at the San Deigo Supercomputer Center through allocation BIO230113 from the Advanced Cyberinfrastructure Coordination Ecosystem: Services & Support (ACCESS) program, which is supported by National Science Foundation grants #2138259, #2138286, #2138307, #2137603, and #2138296.

## Notes

### Competing Interest Statement

The authors have declared no competing interest.

