## Supplemental Figures for "Gene-by-environment interactions and adaptive body size variation in mice from the Americas"

**Figure S1.** Stable isotope analysis (Carbon ($\delta^{13}C$), top; Nitrogen ($\delta^{15}N$), bottom) of hair samples from Suzuki *et al.* [21] of mice sampled from North and South Americas. Population averages can be found in Table S1.

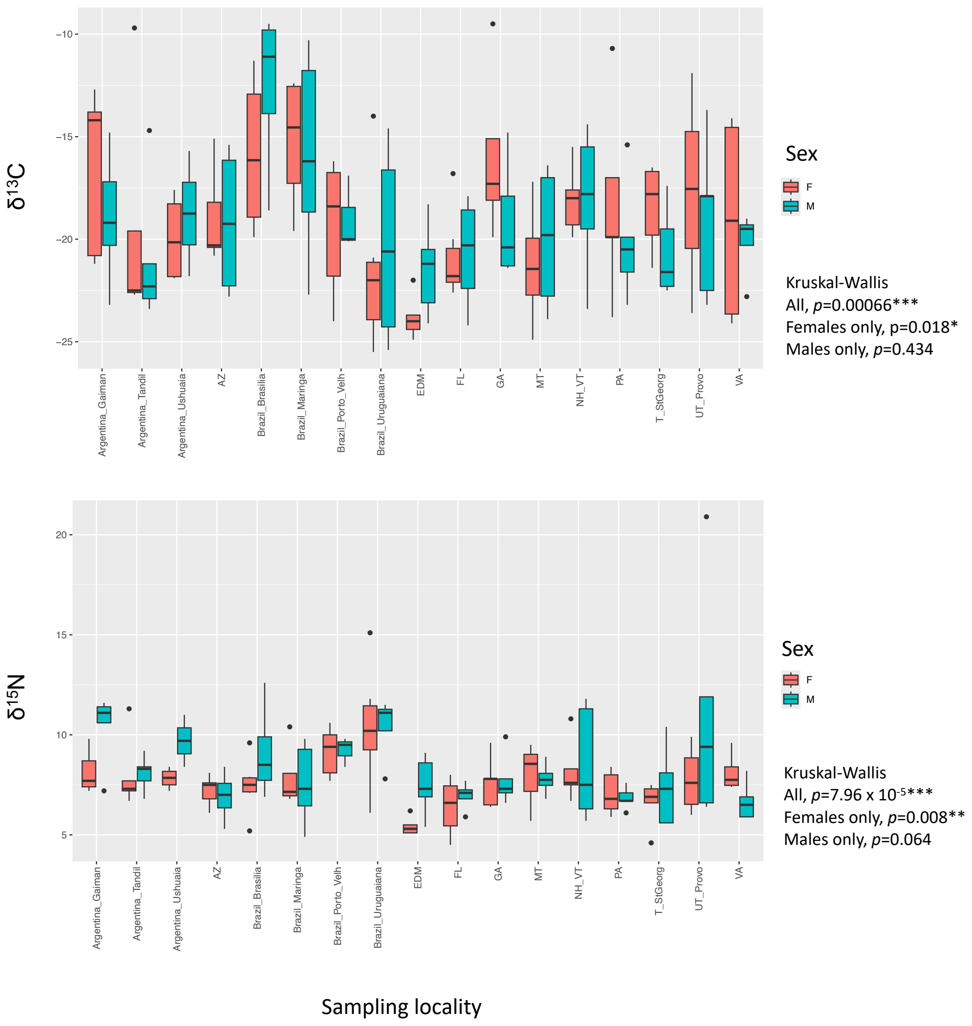

**Figure S2.** Variation in aspects of body size among all strains and diets in: week 12 body weight divided by body length in A) female and B) male mice, growth rate in C) female and D) male mice, BMI in E) female and F) male mice, and body length in F) female and F) male mice.

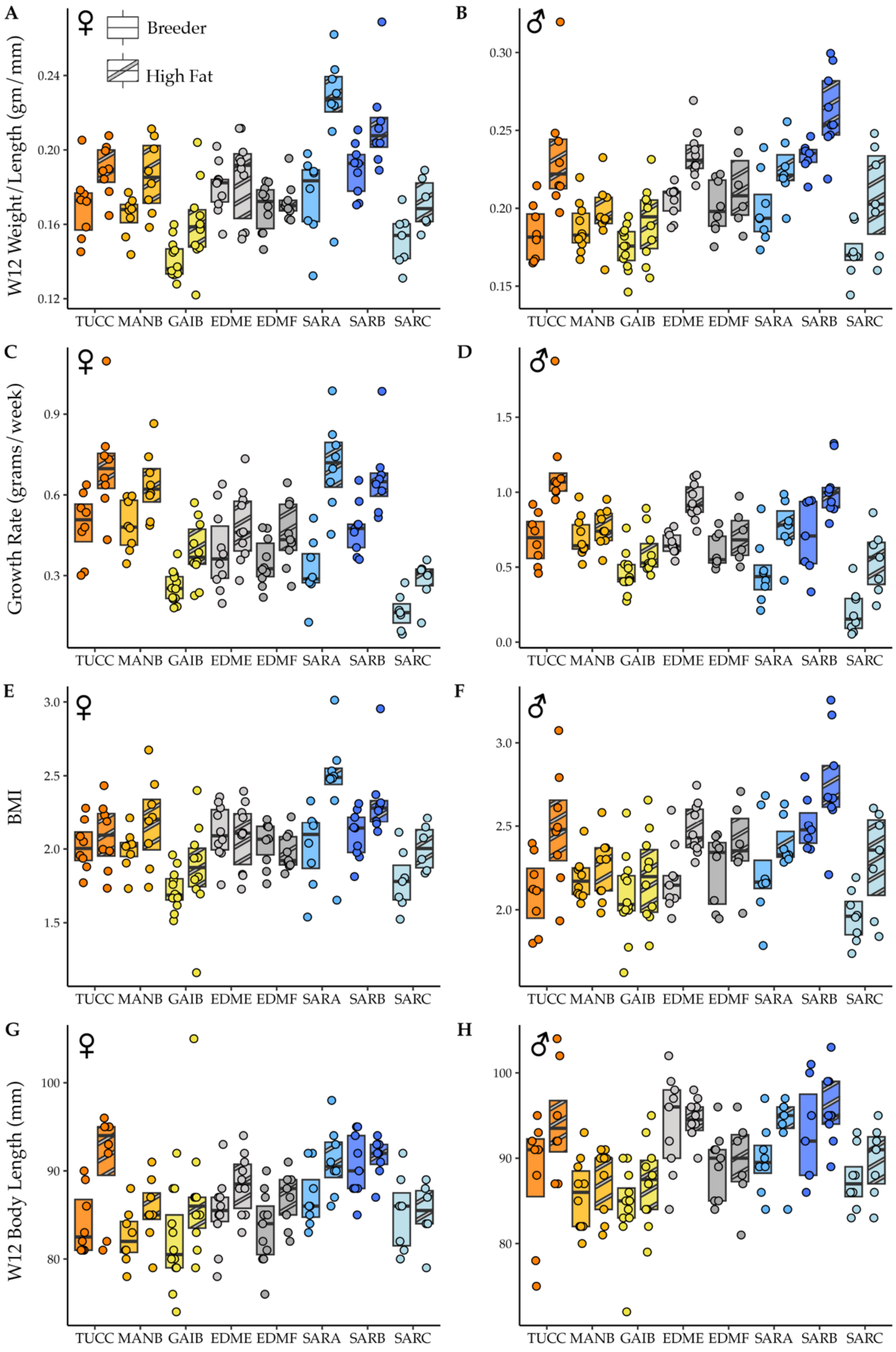

**Figure S3.** Measures of body size on a regular breeder and a high fat diet in the subset of strains selected for transcriptome sequencing. Strain and the interaction between strain and diet contribute to variation in A) body weight and C) body weight divided by body length in female mice. In male mice, strain and diet contributed to variation in B) body weight and D) body weight divided by body length in male mice.

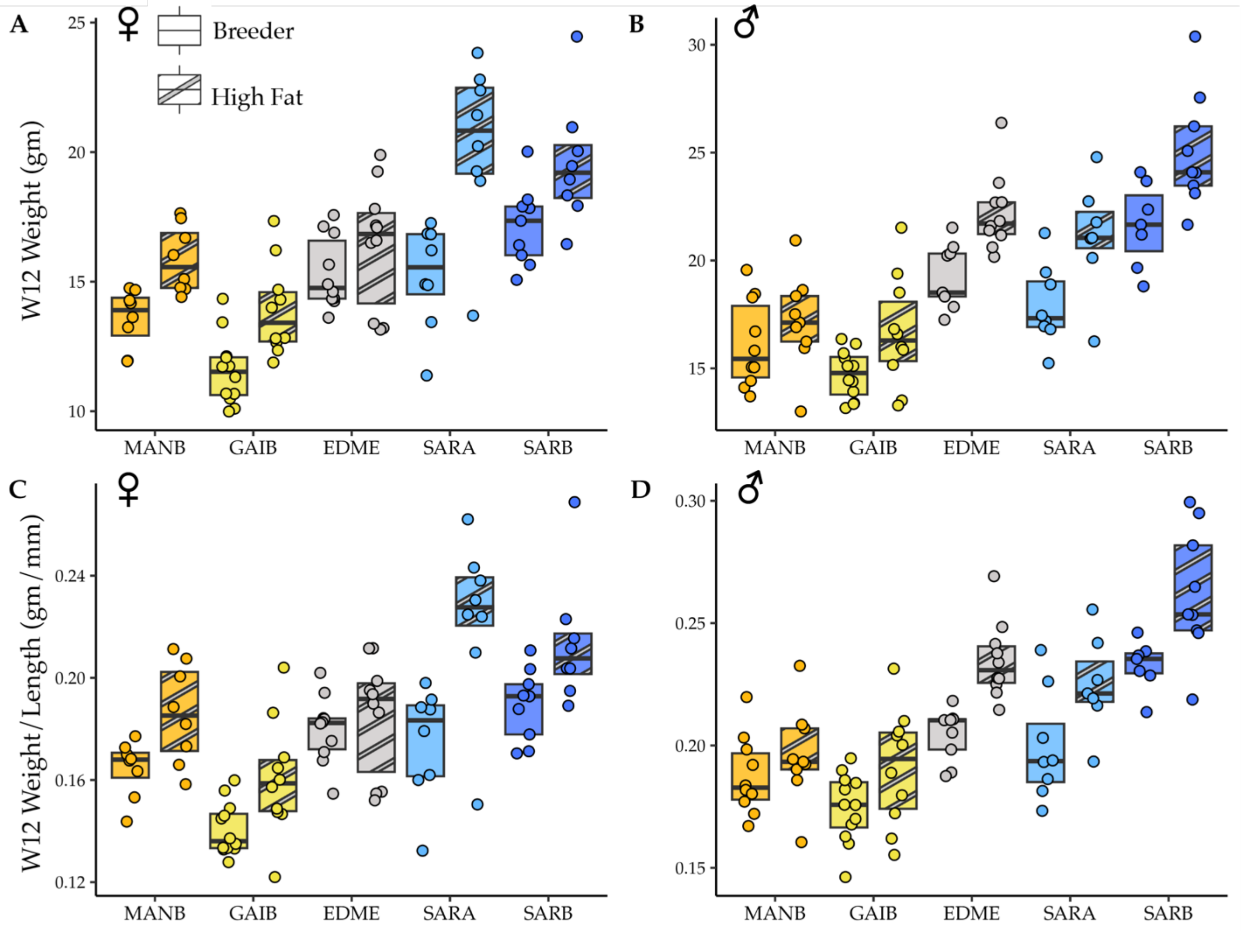

**Figure S4.** PCA of gene expression in females for each strain for high and regular diets (circles =high fat diet, triangle = regular diet).

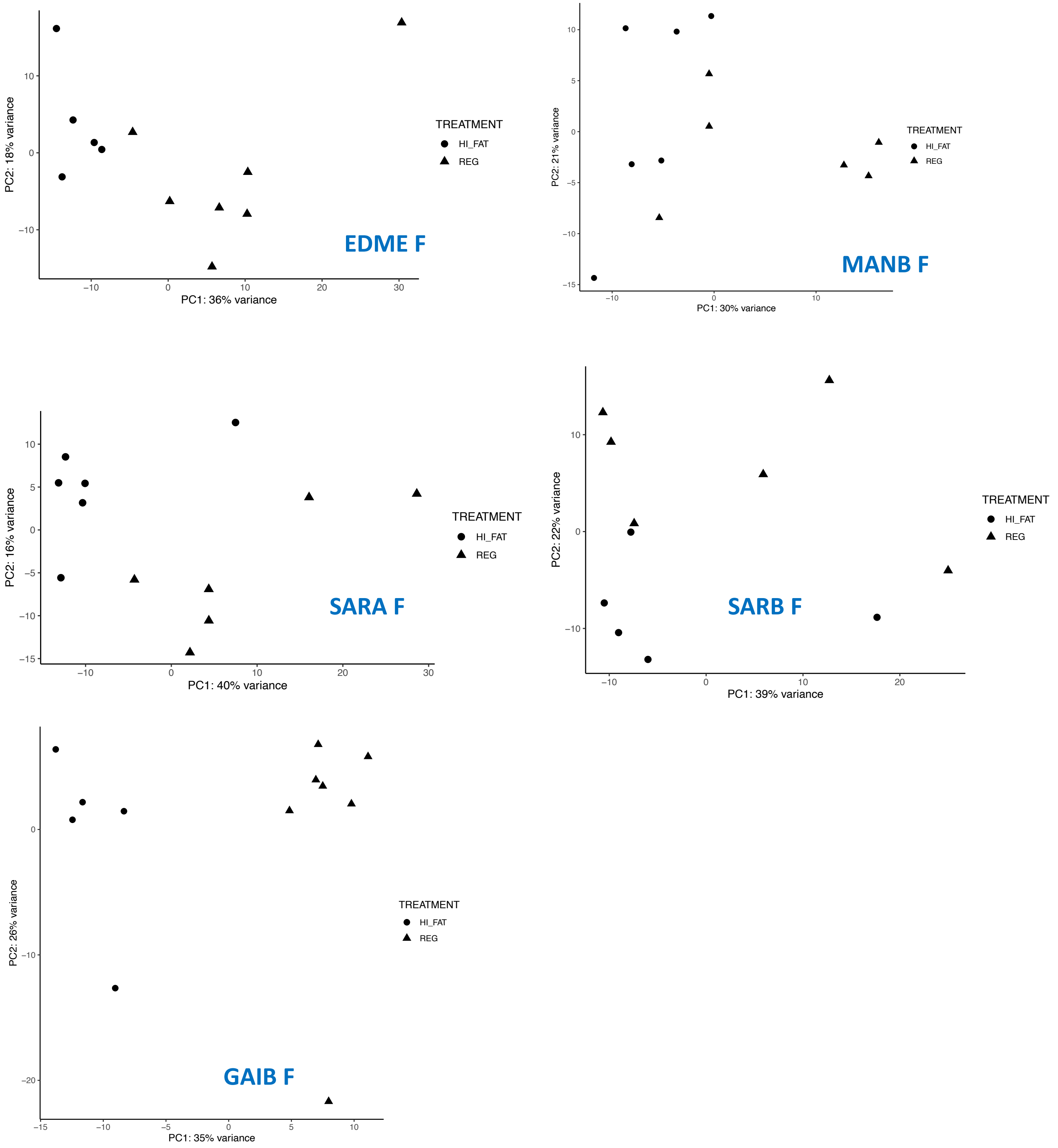

**Figure S5.** PCA of gene expression in males for each strain for high and regular breeder diets (Circles = high fat, triangles = regular).

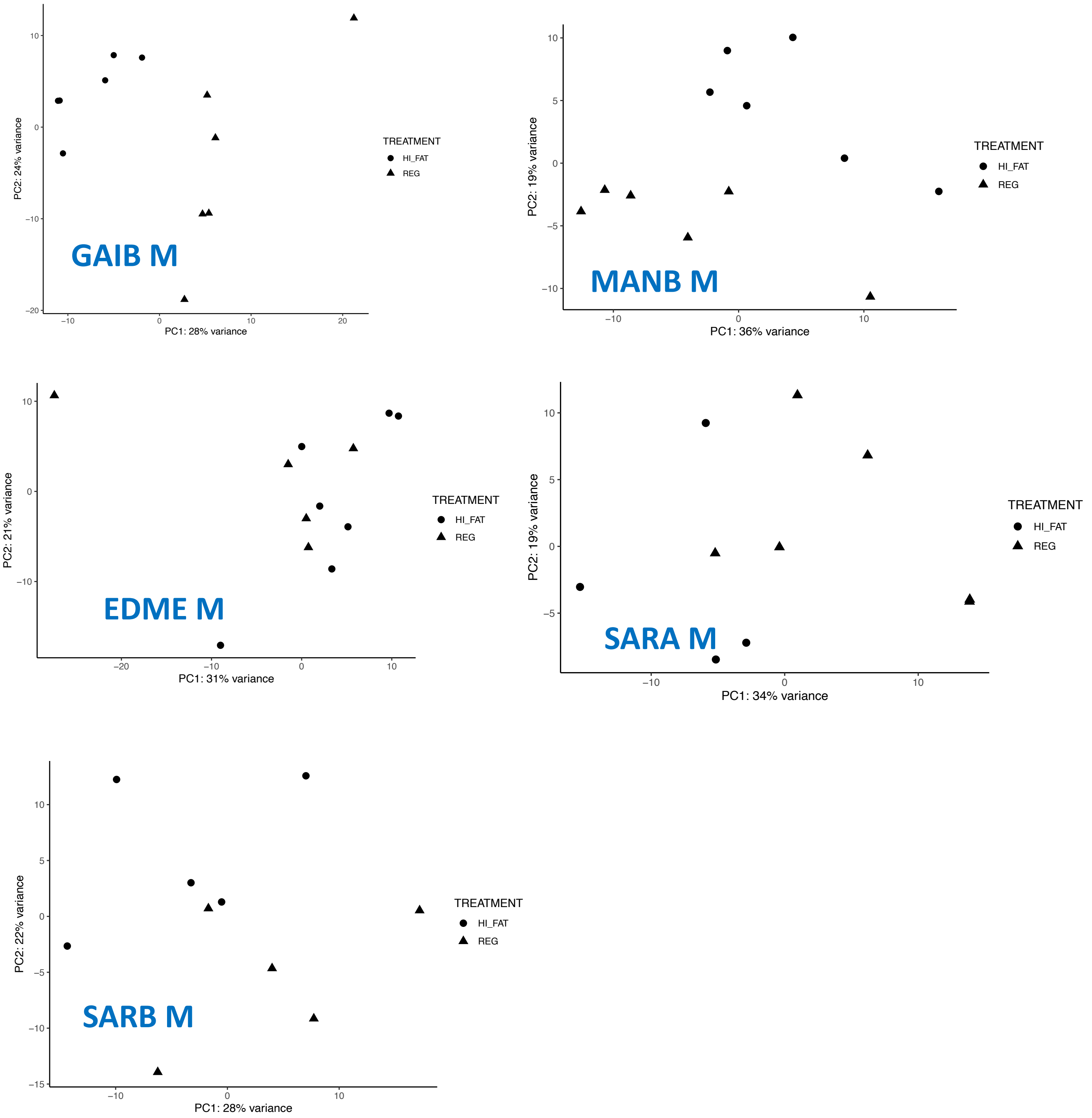

**Figure S6**. Expression of *Gstm2* across strains and diets. Each strain is designated by an acronym (BZ=Manaus, Brazil [MANB], ED=Edmonton, Canada [EDME], FL= Florida, USA [GAIB], NY_1=New York, USA [SARA], NY_2 =New York, USA [SARB]), sex (M=male, F=female), and diet (HI= high fat, REG= regular).

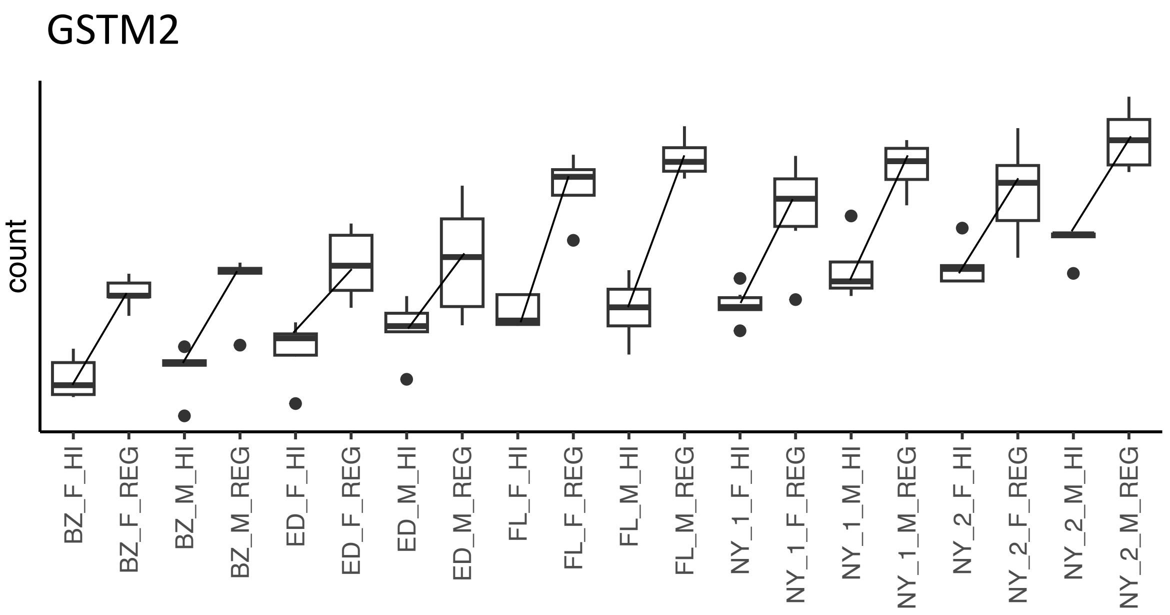

**Figure S7.** UpSet plot showing the overlap of genes with significant transcriptional responses to diet across classical inbred strains.

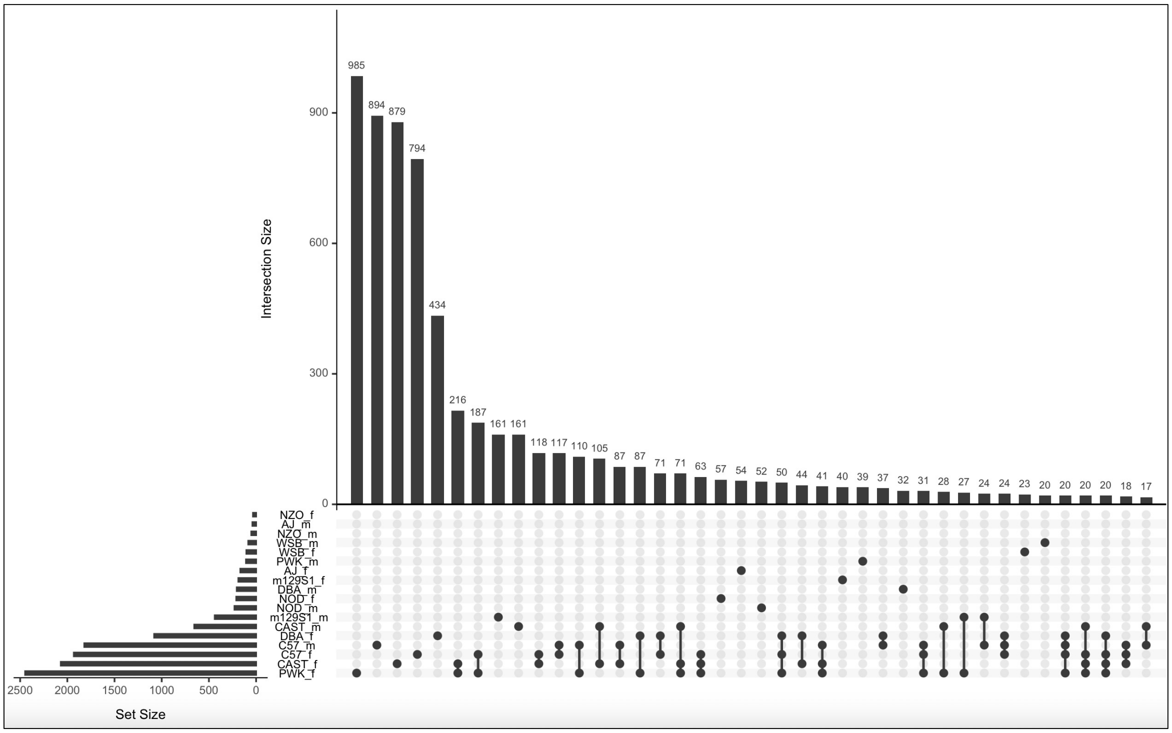

**Figure S8.** UpSet plot showing the overlap of genes with significant transcriptional responses to diet between males of classical inbred lines. Here, NachJ strains are labelled by locality (BZ=MANB, NY_1=SARA, NY_2=SARB, FL=GAIB, ED=EDME).

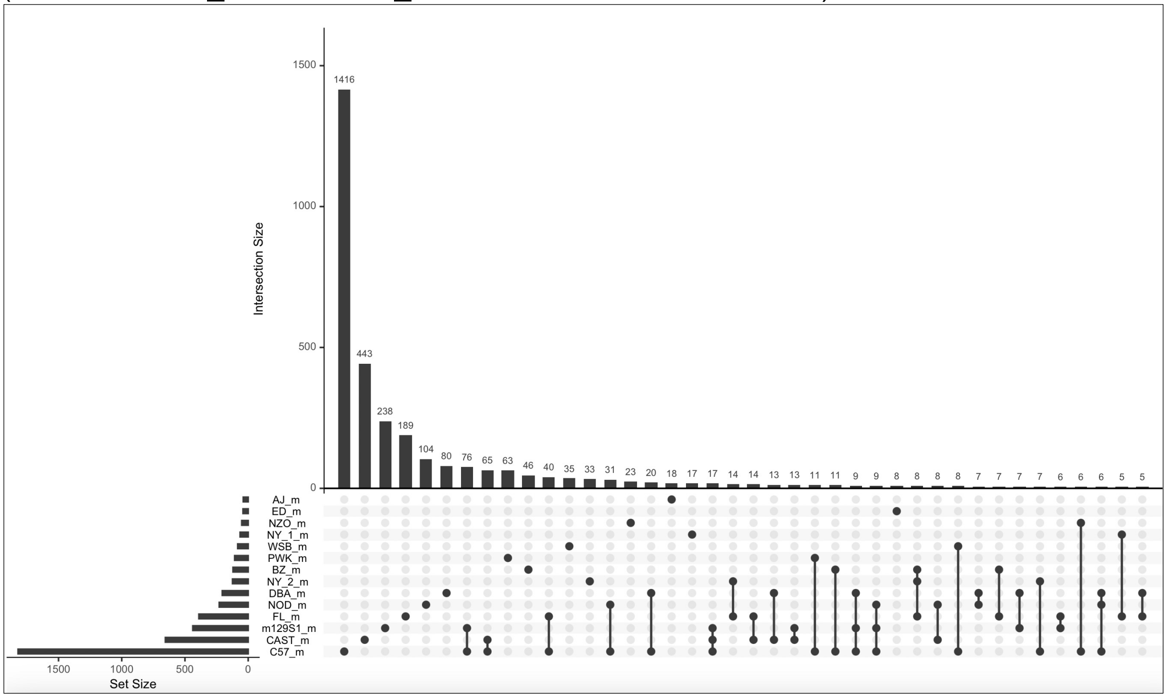

**Figure S9.** UpSet plot showing the overlap of genes with significant transcriptional responses to diet between females of classical inbred lines and NachJ lines. Here, NachJ strains are labelled by locality (BZ=MANB, NY_1=SARA, NY_2=SARB, FL=GAIB, ED=EDME).

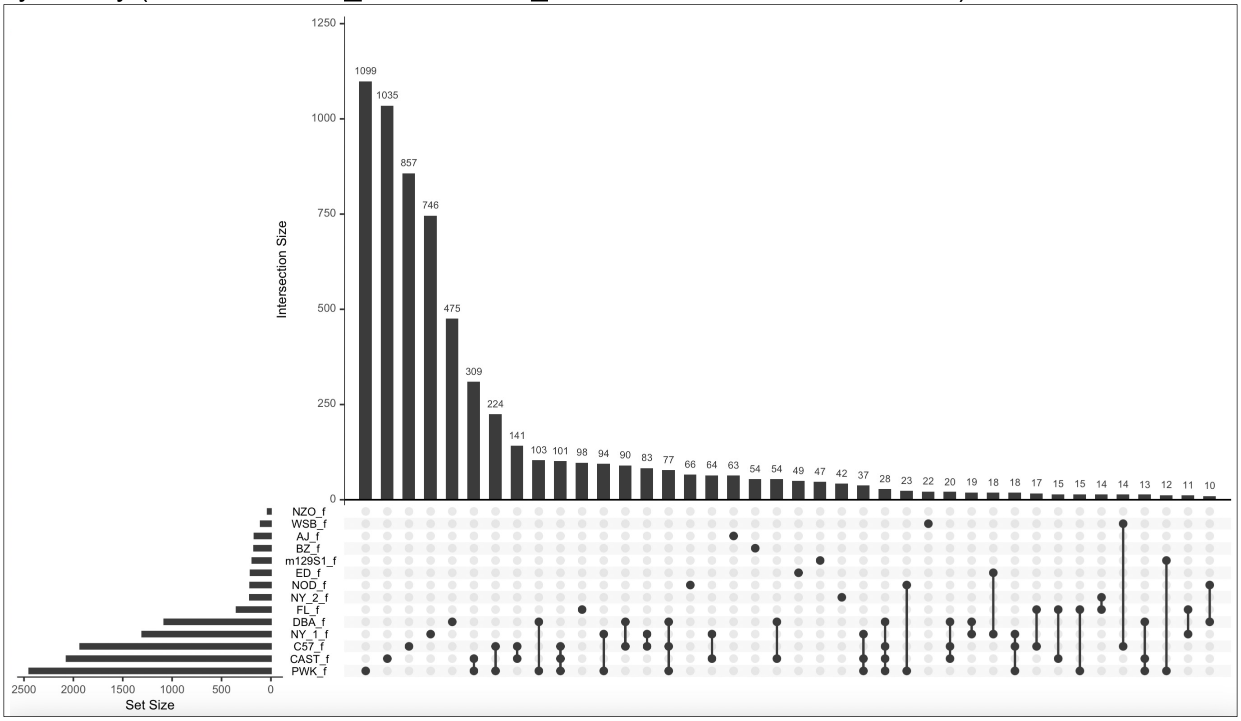

**Figure S10.** Consensus module associations with traits for female mice.

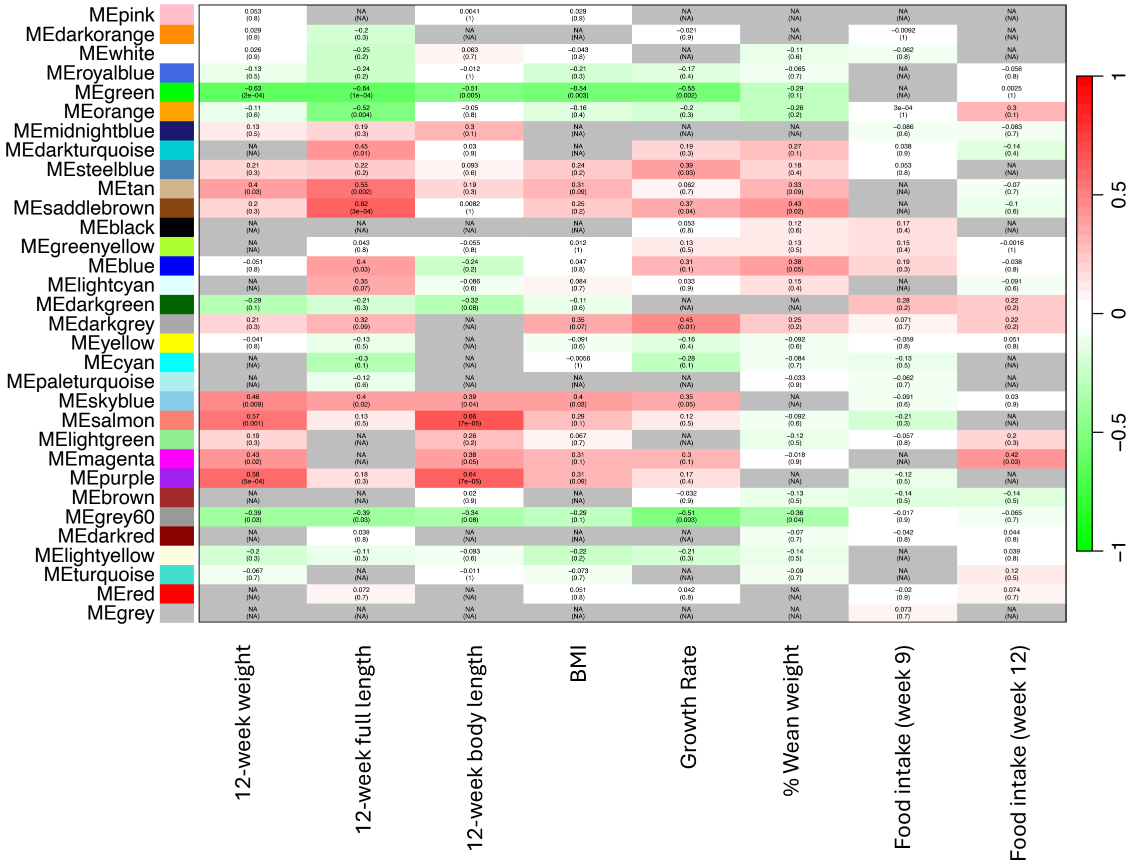

**Figure S11.** Consensus module associations with traits for male mice.

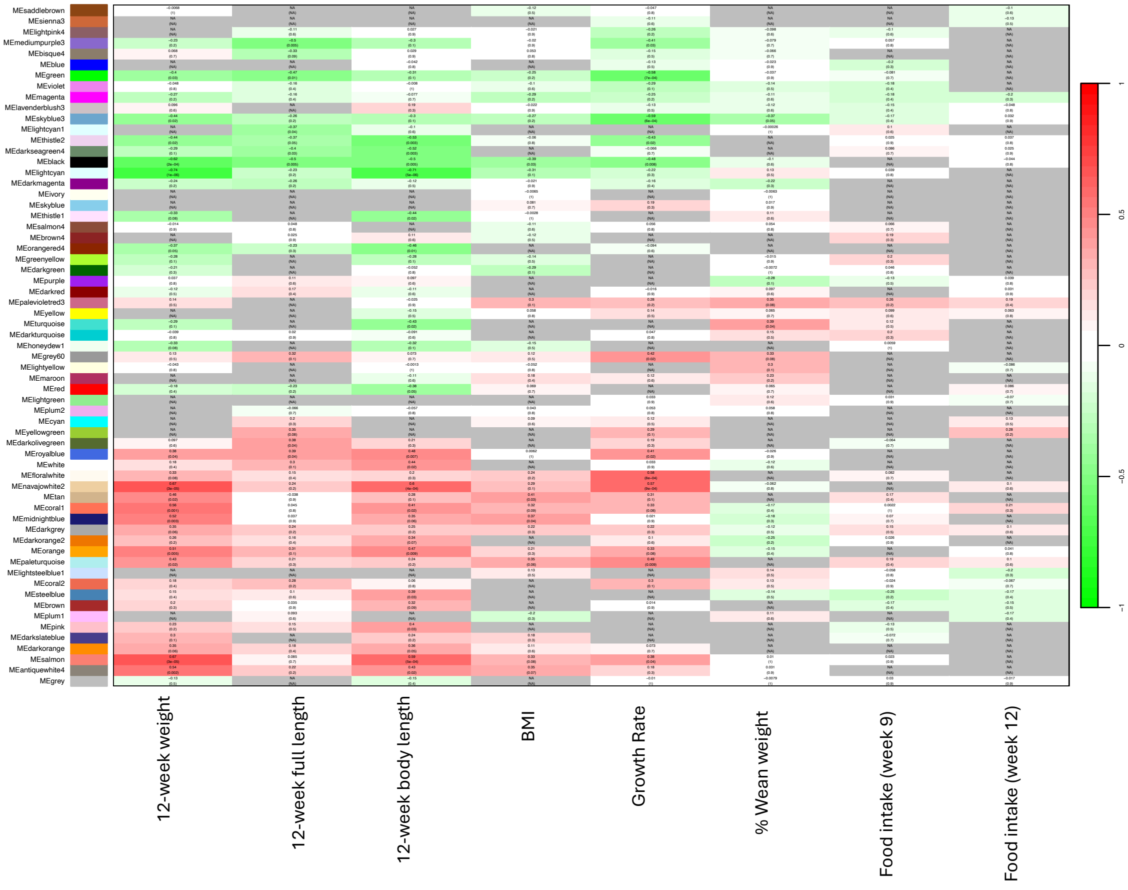

**Figure S12.** Consensus eigengene networks and their differential analysis in female mice. Top panels show dendrograms of consensus module eigengenes (designated by module colors) for regular and high fat diets. The barplot indicates mean preservation of adjacency for each eigengene to other eigengenes. Eigengene networks for the diet sets are shown as heatmaps (labelled “Regular fat diet” and “High fat diet”). Heat maps are colored by level of adjacency (red = high, blue = low). Preservation heatmap (bottom left) shows the preservation of across diets. This plot was created with the “plotEigengeneNetworks” command in WGCNA.

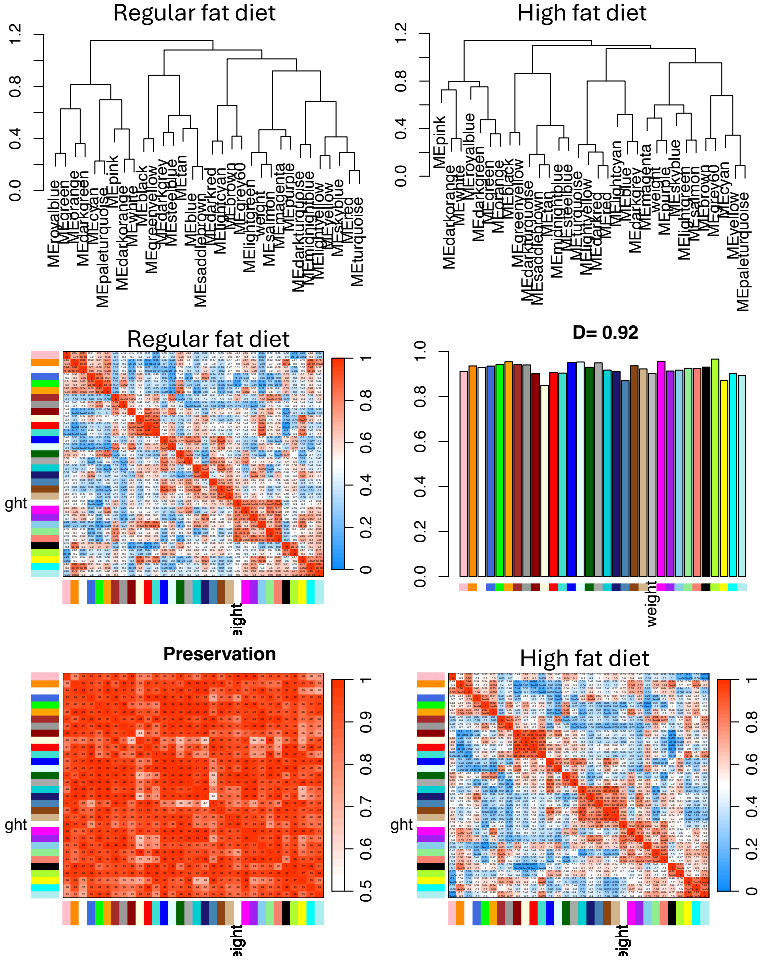

**Figure S13.** Consensus eigengene networks and their differential analysis in male mice. Top panels show dendrograms of consensus module eigengenes (designated by module colors) for regular and high fat diets. The barplot indicates mean preservation of adjacency for each eigengene to other eigengenes. Eigengene networks for the diet sets are shown as heatmaps (labelled “Regular fat diet” and “High fat diet”). Heat maps are colored by level of adjacency (red = high, blue = low). Preservation heatmap (bottom left) shows the preservation of across diets. This plot was created with the “plotEigengeneNetworks” command in WGCNA.

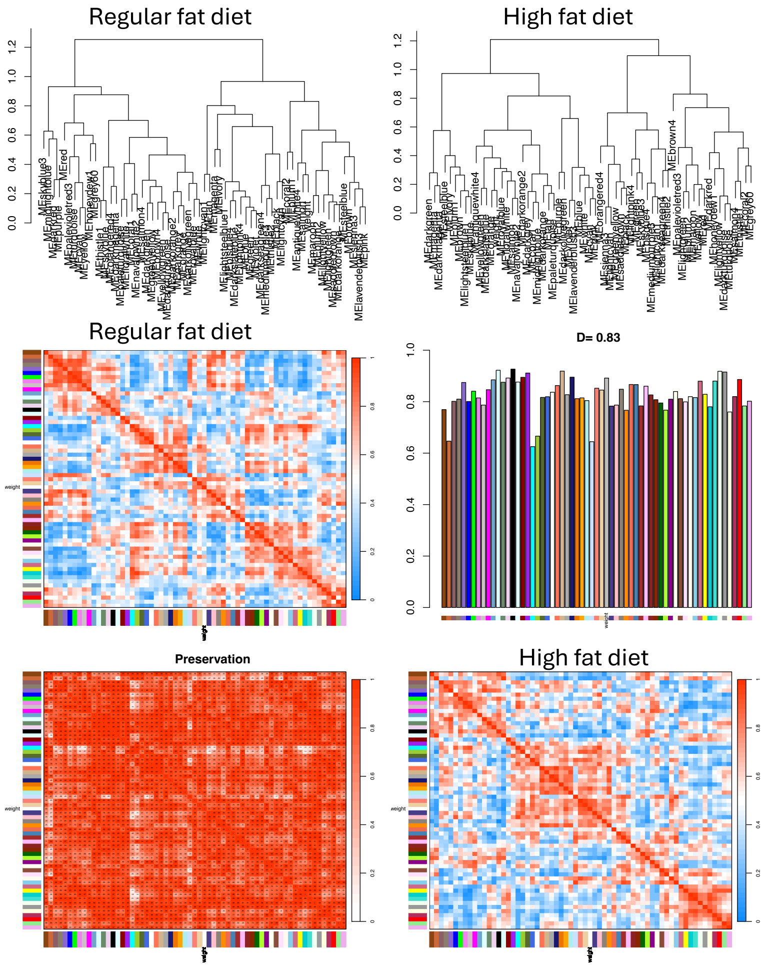

**Figure S14.** Measures of body size on a regular breeder diet or a high fat diet for SARB, MANB, and the cross SARBXMANB. Female Mice A) body weight and C) body weight divided by body length. Male mice B) body weight and D) body weight divided by body length.

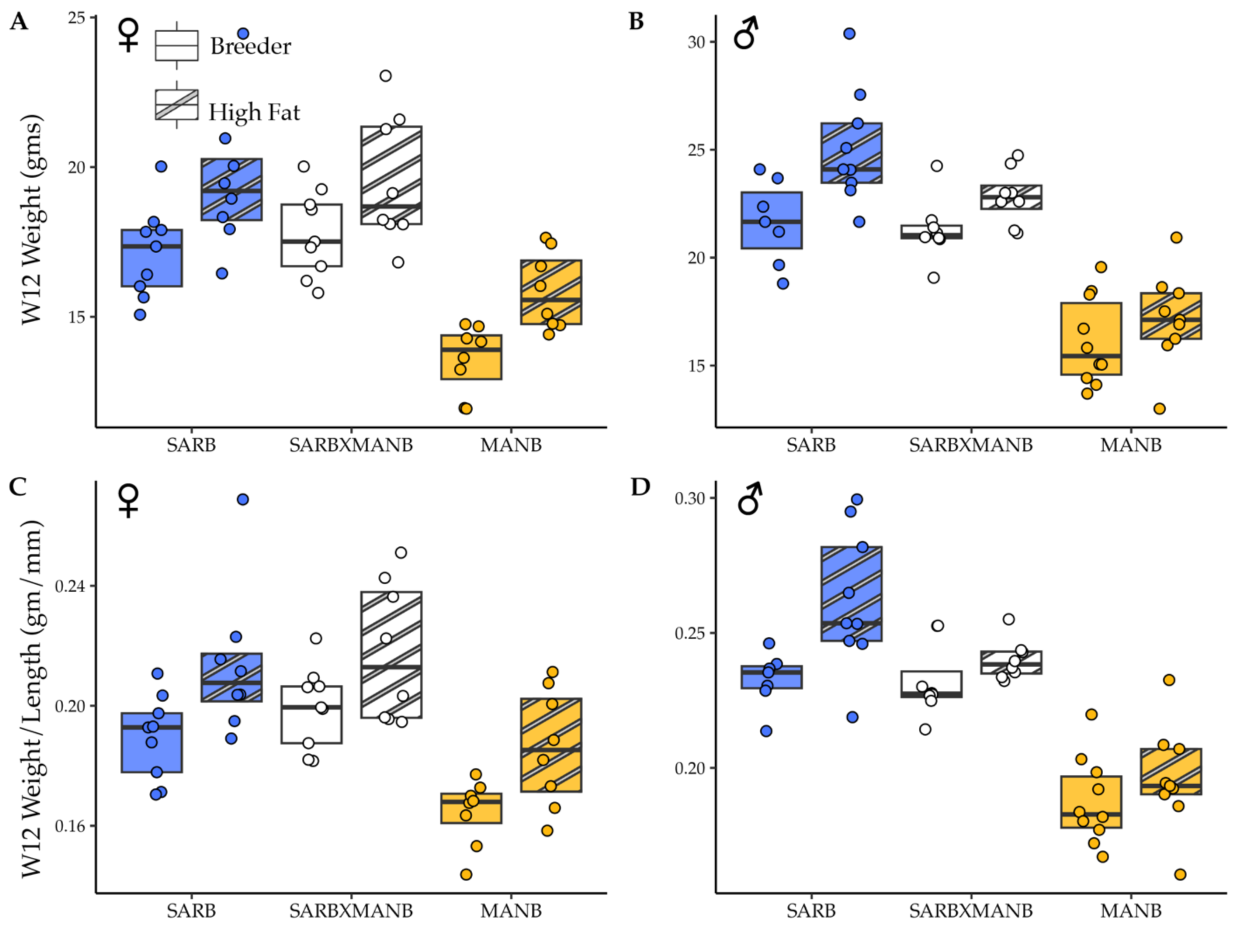

**Figure S15.** Measures of body size on a breeder diet and a high fat diet for GAIB, SARA, and the cross GAIBXSARA. Female Mice A) body weight and C) body weight divided by body length. Male mice B) body weight and D) body weight divided by body length.

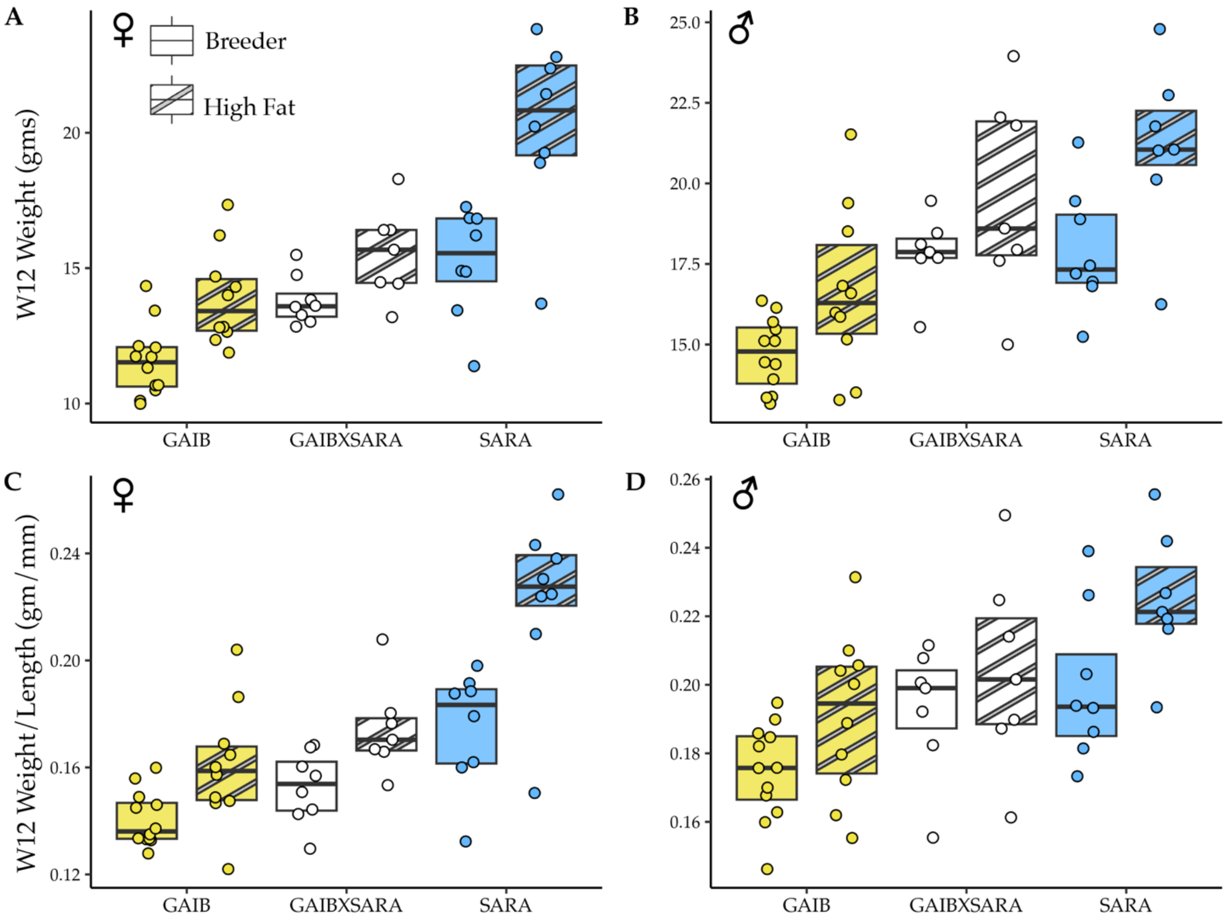

**Figure S16.** Genes with evidence of *cis*-by-diet interactions (red points) based on comparisons between allelic ratios in hybrids on high fat vs. regular fat (breeder) diets.

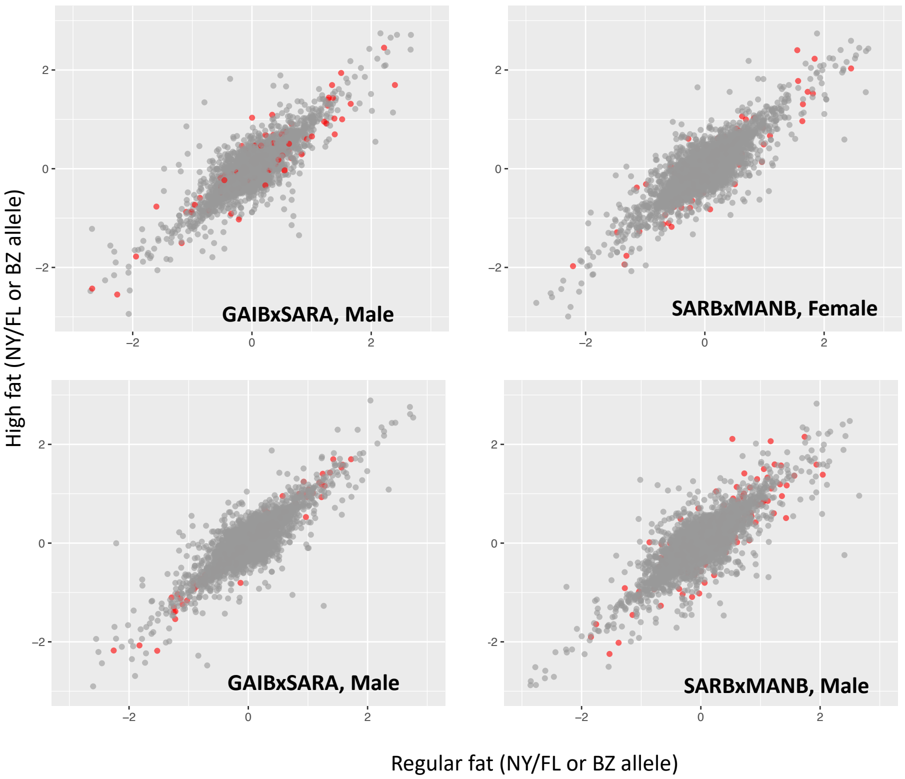
